# Genomic Insights into the Evolution of Parental Care in Weevils

**DOI:** 10.1101/2025.08.27.672529

**Authors:** Sarah Rinke, Peter Biedermann, Martin Schebeck, Mark C. Harrison

## Abstract

Parental care, a key step in the evolution of sociality, has evolved multiple times in insects, yet the molecular mechanisms underlying its emergence remain poorly understood. Weevils (Curculionidae) exhibit diverse parental care behaviours, from nest building to egg and larval attendance, making them an ideal system to investigate genomic changes associated with subsociality. We analysed 13 high-quality weevil genomes, encompassing independent origins of egg and larval attendance, to test two predictions: (1) the sheltering hypothesis, where parental care relaxes selection on traits critical for independent larval survival, and (2) the regulatory hypothesis, where behavioural shifts are driven by changes in transcriptional regulation. Gene family evolution analyses revealed significantly more convergent contractions, particularly in genes linked to transcriptional regulation and enzymatic activity, on branches where larval attendance evolved, consistent with functional gene loss under relaxed selection. Selection analyses identified over 400 genes under relaxed selection, especially those associated with transcriptional regulation and neural plasticity, further supporting hypothesis 1. In contrast, positive selection and intensified selection were rare but enriched for genes regulating gene expression, consistent with hypothesis 2. Together, these results suggest that parental care in weevils drives both simplification of larval traits through relaxed selection and convergent gene loss, and innovation in caregiving behaviours via adaptive changes in gene regulation.

**Significance statement:** Parental care is a pivotal evolutionary innovation, yet its genetic basis in insects remains underexplored. By comparing genomes from weevil species with and without care behaviours, we reveal two key processes shaping the emergence of subsociality: relaxation of selection on genes linked to larval independence and adaptive evolution in genes regulating gene expression. This combination likely reflects reduced demands on protected larvae alongside fine-tuning of parental behaviours. Our findings highlight how simple social systems can evolve through both loss and innovation, offering a comparative framework for understanding social evolution across insects.

## Introduction

Parental care is a social phenotype that is widespread in many vertebrate lineages, especially within mammals and birds. Taking care of offspring increases the offspring’s chances of survival and reproduction while often reducing parental fecundity (Royle et al. 2012). The extent to which a species performs parental care depends strongly on the environment and the type and availability of food (Edward O. Wilson 1975; Douglas W. Tallamy 1984; D. W. Tallamy & Wood 1986). Parental care in insects, where it has evolved in at least 13 orders (Eickwort 1981, as cited in Douglas W. Tallamy 1984), predominantly exists in species feeding on leaves which are exposed to predators and parasites, in species feeding on or living in wood to inoculate their offspring with symbionts, as well as in insects feeding on dung, which is an ephemeral and highly competitive resource (Royle et al. 2012). In beetles, most families in which parental care has evolved feed on these food types (Biedermann & Nuotclà 2020), for instance, leaf beetles (Chrysomelidae), dung beetles (Scarabaeidae), and wood-boring weevils (within Curculionidae).

Insects display a broad range of social phenotypes from parental care, also called subsociality, up to the most complex social phenotype, eusociality, which evolved in three insect orders: ants, bees and wasps within Hymenoptera, termites within Blattodea, and at least one species of ambrosia beetle (*Austroplatypus incompertus*) within Coleoptera. Eusociality encompasses the presence of alloparental care, overlapping generations and division of reproductive labour into reproductive and sterile castes (E. O. Wilson 1971). Studies of less socially complex lineages, such as subsocial species without castes or social species with monomorphic, flexible castes, are important for understanding the evolution of eusociality (Linksvayer 2010; Shuker & Simmons 2014; Kronauer & Libbrecht 2018). However, research on the molecular evolution of sociality in insects has traditionally focused on eusocial species, especially within Hymenoptera (e.g. Kapheim et al. 2015; Simola et al. 2013; Shell et al. 2021; Jones et al. 2023) and less frequently in Blattodea (Harrison et al. 2018; Ewart et al. 2024).

It has been proposed that at different levels of social complexity in insects, different molecular mechanisms may play a role in the evolution of social traits (Rehan & Toth 2015). These expectations include, for example, changes in gene expression patterns that affect behavioural shifts in simple societies, while greater adaptational changes in gene repertoire are expected to be involved in the evolution of divergent caste phenotypes (Rehan & Toth 2015; Linksvayer & Wade 2005). Most of these expectations of gene family size evolution, adaptive protein evolution and evolutionary changes in transcriptional regulation have been confirmed with comparative genomic and transcriptomic analyses as being important for the evolution of increasing social complexity within eusocial taxa, such as bees, ants, wasps (Hymenoptera), and termites (Blattodea) (Kapheim et al. 2015; Shell et al. 2021; Harrison et al. 2018; Simola et al. 2013). However, subsocial species are mostly missing in these studies so that mechanisms involved in the evolution of simple social behaviour such as parental care could not be distinguished from those associated with the evolution of more complex social phenotypes (reviewed in Mikhailova et al. 2024). There are two main hypotheses regarding the molecular mechanisms related to the evolution of parental care: Parental care itself is suggested to lead to less intense selection due to sheltering of the offspring. In this scenario, parents compensate for non-optimised traits of their offspring, thus increasing chances of survival and decreasing selective pressure on the involved traits. This can lead to the accumulation of slightly deleterious gene variants and a dependency on parental care as observed in *Nicrophorus vespilloides* (Mashoodh et al. 2023; Pascoal et al. 2023). A second hypothesis proposes that adaptive changes in gene expression can evolve relatively easily through modifications in the regulation of transcription and translation, often resulting in altered timing or tissue-specific gene expression (Romero et al. 2012). This shift in gene expression is often regarded as an early step in the evolution of sociality and may play a role in the transition from solitary life style to parental care behaviour (Rehan & Toth 2015). Here, we aim to test which evolutionary mechanisms occur at the emergence of subsociality in beetles. More specifically, we investigate if there is evidence for genomic changes in transcriptional regulation mechanisms and a relaxation of selection, which are expected to occur at the emergence of subsociality. Additionally we investigate evidence for adaptive evolution, which is proposed to play a role in the evolution of higher social complexity.

Although parental care evolved in numerous beetle families, the highest number of evolutionary origins of parental care exist in the “true weevils” (Curculionidae, six origins) (Biedermann & Nuotclà 2020). Weevils feed on a great variety of plant parts and fungal tissues and are often categorized based on larval feeding modes (Kirkendall et al. 2015). Sociality level often, but not always, correlates with feeding pattern. Generally, free living weevils have a less complex social lifestyle and feed on a variety of plant parts, like seeds (Spermatophagy), fleshy (not woody) plant tissues (Herbiphagy), some bark beetles feed on phloem, while ambrosia beetles have a more complex social lifestyle and live in woody tissues where they cultivate fungi (Xylomycetophagy) (Kirkendall et al. 2015). However, *Hypothenemus hampei* larvae, for example, feed on seeds, while their parental care behaviour is more complex than in other species with a similar feeding mode. While complex social systems, like (facultative) eusociality, evolved in bark and ambrosia beetles, which is generally not the case in most free-living weevils (Biedermann & Nuotclà 2020). Bark and ambrosia beetle species, cover the sociality spectrum from gregariousness, via parental care, all the way to eusocial societies (Costa 2006). Furthermore, due to their pest behaviour a large number of published weevil genomes are publicly available allowing evolutionary analyses into the emergence of social phenotypes in this group.

Parental care behaviour in weevils is less well explored than, e.g. in carrion beetles, as most species with parental care have a cryptic lifestyle, although more detailed observations are becoming available (Biedermann & Nuotclà 2020). Nevertheless, a simple classification into no care, nest building, egg attendance and larval attendance is possible for most weevil species with published genomes (Kirkendall et al. 2015). Nest building is the simplest form of parental care and can be defined by concealing the eggs from their surroundings by placing them in or under plant materials to prevent predation (Royle et al. 2012). Free living weevils do not build complex nests, but they lay their eggs in well-protected environments for their offspring and often cover the eggs, e.g. by rolling leaves around them. Nest building is relatively simple in species like the *Ips* bark beetles that mate in the phloem and lay individual eggs that they cover with boring dust, but more advanced in the fungus farming ambrosia beetles that live in groups (Kirkendall et al. 2015). In some species, parents stay with their eggs at the oviposition site until the larvae hatch, to protect and groom the eggs. This behaviour is called pre-hatching care or egg attendance. Fewer species perform larval attendance, or post-hatching care, where parents attend to their hatched larvae to protect and groom them (Royle et al. 2012).

In this study, we carried out evolutionary analyses on 13 weevil species with published genomes, covering different parental care phenotypes, namely nest building, egg attendance and larval attendance (Fig. 1). Larval attendance, which represents the most complex social phenotype in this study, has emerged twice within the species we included, allowing us to investigate convergent evolutionary mechanisms involved in the evolution of parental care. We employed selection and gene family size evolution analyses to investigate the role of sheltering and transcriptional regulation in the evolution of parental care in weevils.

**Figure 1:**
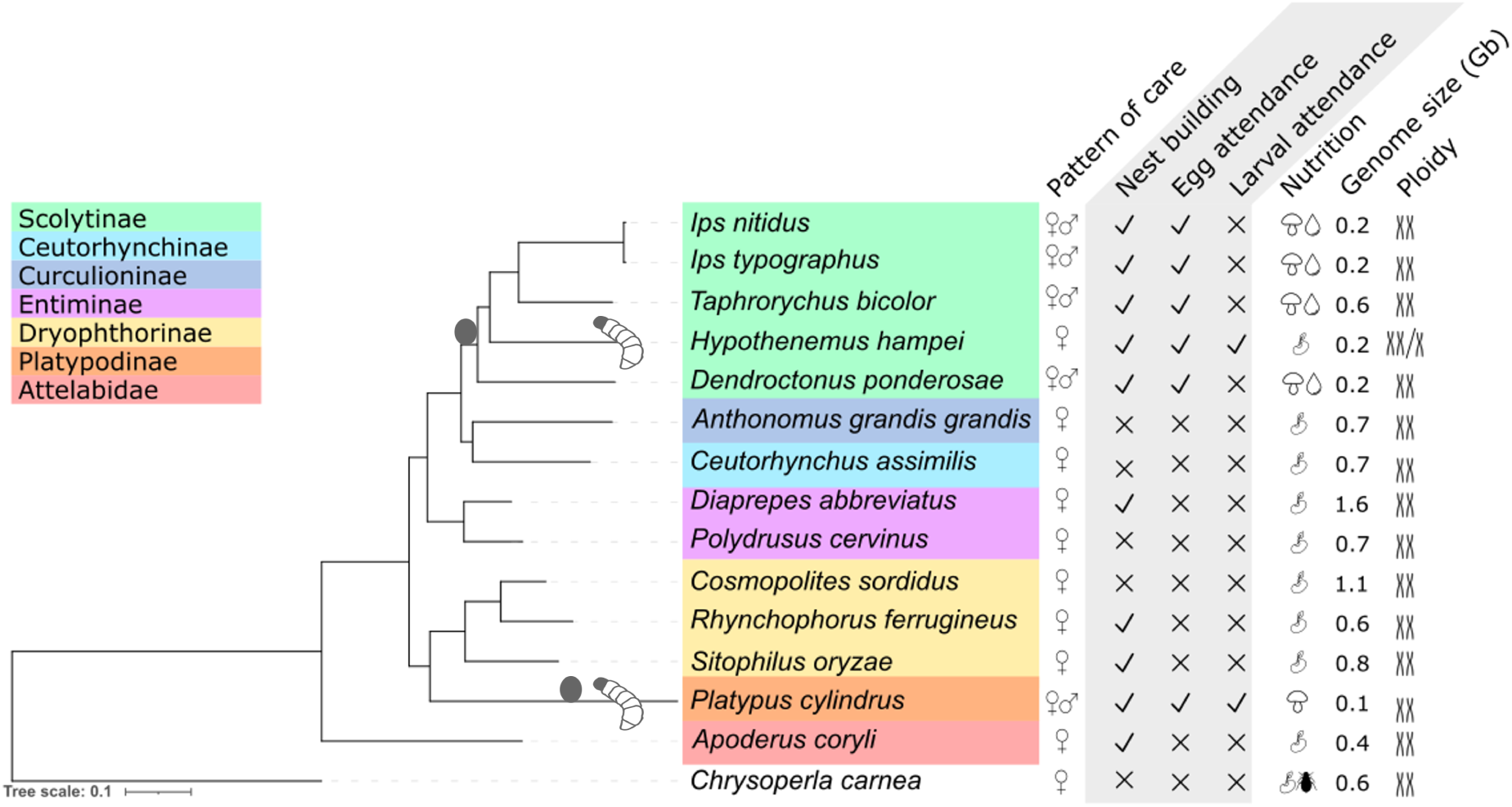
Phylogenetic tree of the studied species with information on parental care, nutritional and genomic traits. The colours indicate the weevil subfamily, eggs indicates the evolution of egg attendance, while the larvae indicate the evolution of larval attendance. Pattern of care is either uniparental female of biparental care. Form of care: all beetles in this tree select the oviposition site (not shown in figure). Nest building, egg attendance and larval attendance are shown as present (✓) or absent (x). The different nutrition types are indicted as follows: fungus and phloem, plant material, fungus, plant material, and aphids. Genome sizes are shown in gigabases. Ploidy is indicated as either diploid or haplodiploid.

## Results

### Data set

We include genomes of 13 species from the weevil family (Curculionidae), along with two outgroups, *Apoderus coryli*, belonging to the leaf-rolling weevils (Attelabidae), and *Chrysoperla carnea*, belonging to the Chrysopidae family (Fig. 1). The analysed weevils show uniparental female care or biparental care, spanning a variety of levels of parental care traits. Nest building, which we define as depositing eggs in a well-protected environment and potentially covering the eggs, is present in ten of the analysed species. Within our data set, egg attendance, the grooming and protection of the eggs, is only performed by members of the Scolytinae tribe and *Platypus cylindrus* (Platypodinae). Adults of only two species, *Hypothenemus hampei* and *P. cylindrus*, attend to their offspring after hatching. The food sources range from different plant tissues in most species to fungus farming in ambrosia beetles. Genome sizes vary between 0.1 gigabases in *P. cylindrus* and 1.6 gigabases in *Diaprepes abbreviatus*. All species are diploid, except for *Hypothenemus hampei*, which is haplodiploid.

The focal branches in this study are those on which egg attendance and larval attendance evolved (eggs/larvae in Fig. 1). Parental care evolved at least twice within the weevil family. In the case of Scolytinae, egg attendance evolved in the ancestor of all Scolytinae species used in this study and larval attendance evolved in *H. hampei*. For Platypodinae, both, egg and larval attendance evolved on the branch leading up to *P. cylindrus* (Fig. 1).

The publicly available genomes obtained from NCBI had a BUSCO (Manni et al. 2021) completeness score of ≥ 97.0%, indicating high quality assemblies. All published genomes were re-annotated to ensure uniform proteome quality for downstream analyses. The quality of the re-annotated proteomes was tested with both, BUSCO and DOGMA (Dohmen et al. 2016), yielding very high levels of completeness of ≥ 91.6% (Supplementary Table S1).

### Gene family size evolution

We analysed gene family size changes across the phylogenetic tree to see if expansions and contractions of gene families correlate with differences in parental care phenotypes, specifically egg and larval attendance. For this we used OrthoFinder (Emms & Kelly 2019) to cluster protein sequences into orthologous groups (∼ gene families). CAFE5 (Mendes et al. 2021) was then used to reconstruct sizes of gene families for ancestral nodes and then infer changes in gene family size. Generally, more contractions (purple) than expansions (turquoise) were observed (Fig. 2), even more so in the extant species compared to the ancestral branches. There were no systematic changes in gene family size changes on the branches leading up to species with egg or larval attendance.

**Figure 2:**
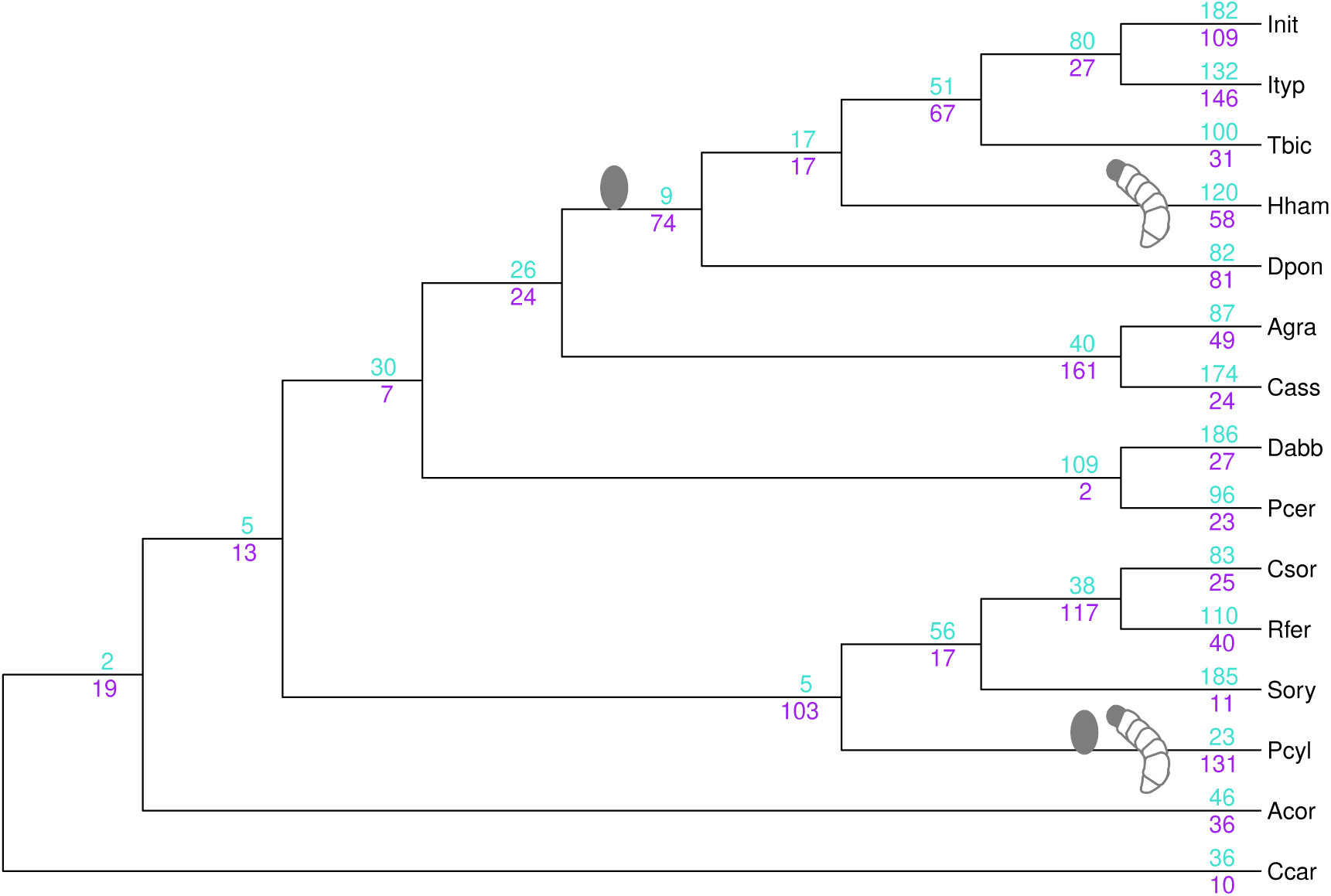
Gene family size changes across the phylogenetic tree. CAFE5 was used to calculate expansions and contractions, numbers indicate the significant changes. All turquoise numbers (above branches) indicate gene family size expansions, while purple numbers (below branches) show contractions. Eggs denote where egg attendance evolved, larvae denote where larval attendance evolved. For full species names please check Table S1.

Next, we focused on gene families that expanded or contracted on the three branches of interest where egg or larval attendance evolved, namely ancestor of Scolytinae species, *P. cylindrus* and *H. hampei* (eggs and larvae in Fig. 1). On the branch leading up to all Scolytinae species, 9 gene families were expanded, while 74 were contracted. In *H. hampei*, there were 120 gene family expansions and 58 contractions, while in *P. cylindrus*, 23 gene families were expanded and 131 were contracted (Fig. 2).

To infer convergent evolution of gene families along with the emergence of egg and larval attendance, we considered numbers of gene families significantly expanded or contracted on more than one of the three branches (Fig. 3). Next we compared these values to expectations based on shared expanded and contracted gene families among all combinations of two branches across the tree (Fig. 3). Overall, the number of convergent expansions between two branches where egg and/or larval attendance evolved (3, 2, 0) was lower than expected based on the numbers of convergent expansions found in all combinations of two branches (median: 4; Fig. 3). The number of convergent contractions on pairs of these focal branches (32, 16, 4), on the other hand, was higher than expected, with a median number of convergent contractions between any two branches in the tree equal to 2 (Fig. 3). A superexact test showed the convergent contraction of 32 gene families on the *P. cylindrus* and *H. hampei* branches, where larval attendance evolved, to be significant (expected: 8.88; p=2.066E-13; Fig. 4). Functions of these convergently contracted gene families could be related to the evolution of larval attendance, which emerged on each branch. Functional annotation of these gene families reveals a variety of functions (Table S2) involved in gene expression regulation, such as endonuclease, ribonuclease H protein, zinc finger protein, histone 2A, acyl-Coenzyme A oxidase, AAA domain containing protein. Also many gene families with enzymatic activity were found, specifically peptidase S1 family serine endopeptidase, choline dehydrogenase, lipase, serine protease, alcohol-forming fatty acyl-CoA reductase, acyl-Coenzyme A oxidase. Other functions like transport of molecules, MreB/Mbl protein / Heat shock protein, thaumatin family were also enriched in these contractions. No GO-terms were enriched within these 32 gene families underlining the diverse functions of these convergently contracted genes.

**Figure 3:**
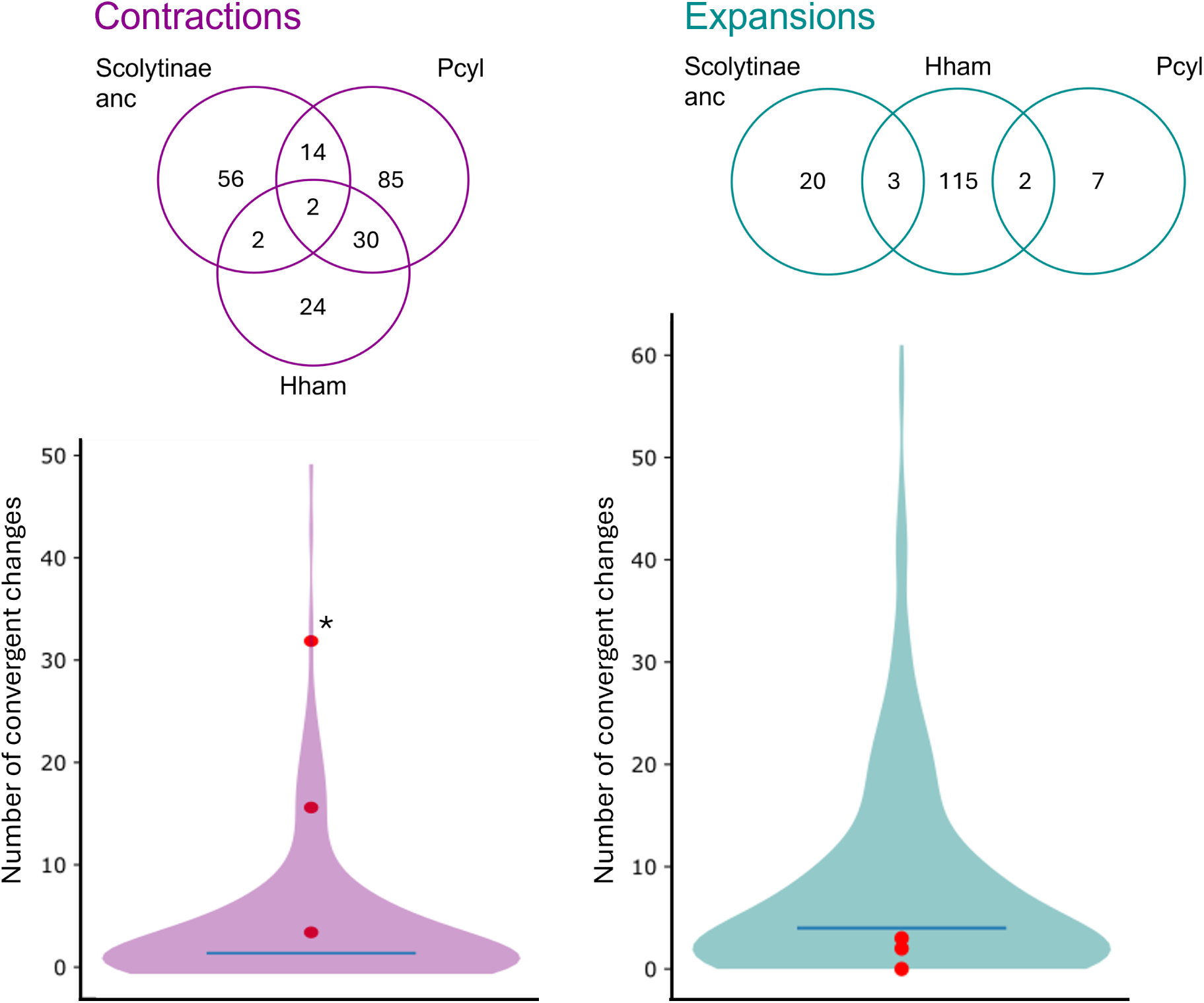
Convergent contractions (purple) and expansions (turquoise). Top: Venn diagrams of convergent contractions (purple) and expansions (turquoise) on the three branches where egg and/or larval attendance evolved. Bottom: Number of observed convergent gene family size changes (purple - contractions, turquoise - expansions) on two branches of interest (red dots) compared to the distribution of number of gene families convergently contracted/expanded in all possible combinations of two branches across the tree. The blue line indicates the median number of convergent expansions in all possible combinations of two branches. Asterisk indicates statistical significance in superexacttest (Fig. 4). For full species names please check Table S1.

**Figure 4:**
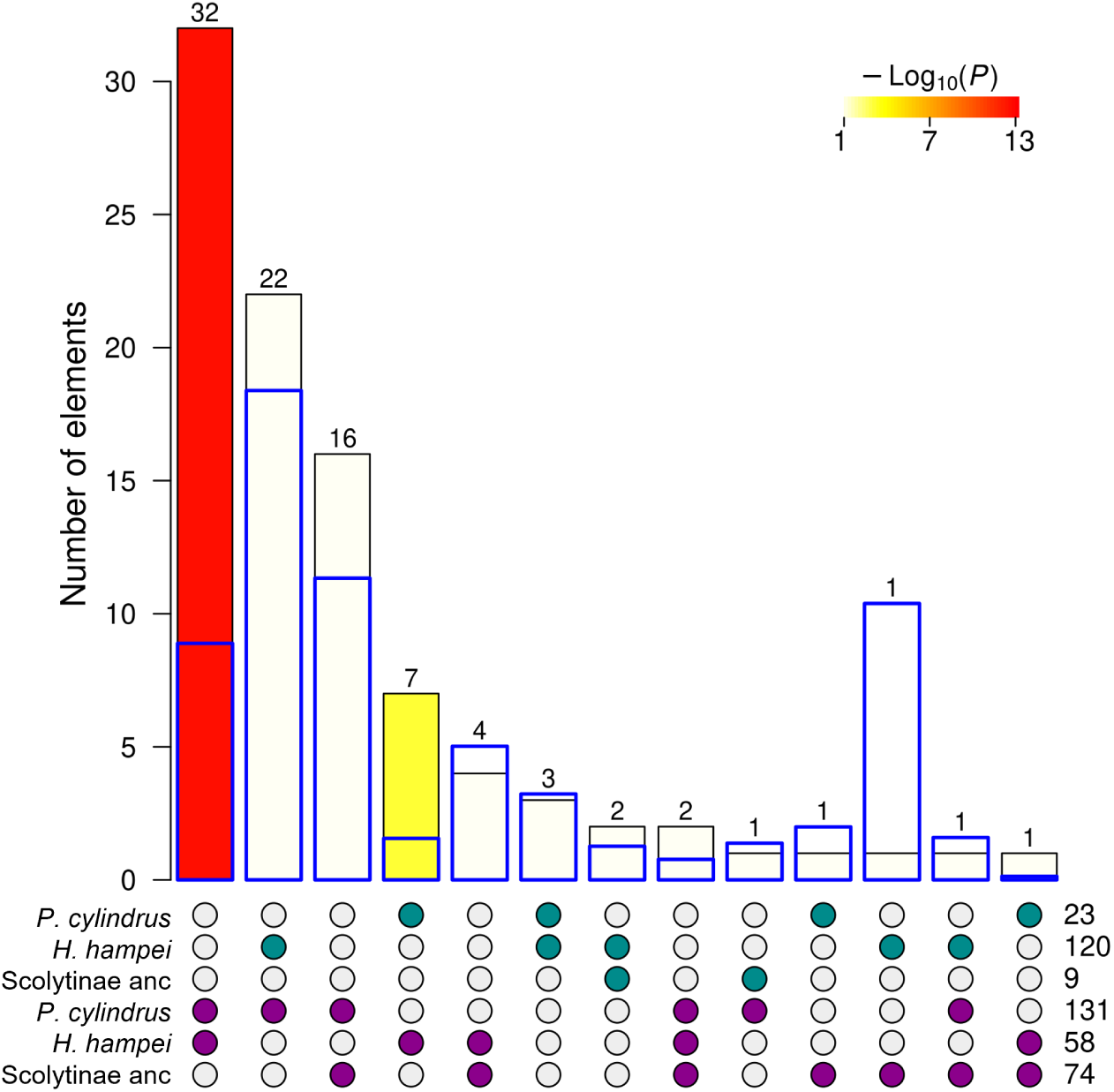
Superexact test of overlaps of gene family expansions and contractions on the three branches of interest, namely ancestor of Scolytinae, *P. cylindrus* and *H. hampei*. Filled circles at the bottom of the figure indicate, which expansions (turquoise) and contractions (purple) are compared. The total number of expansions/contractions for each branch of interest is shown in the bottom right. Blue boxes indicate the expected overlap calculated from the superexact test. Yellow and red shading indicate the p-value is significant at p≤ 0.05 level.

The two gene families that were convergently contracted on all three branches of interest indicate a common set of genes involved in the emergence of egg and larval attendance. The functions of these genes include lipase. One of the three gene families that are convergently expanded on *P. cylindrus* and *H. hampei* is Cytochrome P450, which plays an important role in detoxification in weevils (Blomquist 2021). However, Cytochrome P450s are significantly expanded and contracted on multiple branches in our dataset, including expansions in *Diaprepes abbreviatus* and *Ips nitidus*, both species without larval attendance (Fig. S1).

### Selection

To understand whether the emergence of subsociality has an effect on selection strength, we analysed dN/dS on 2056 single copy orthologues across the phylogenetic tree using codeml from the paml software package (Yang 2007).

We observed strong purifying selection (median dN/dS < 0.05) across the entire phylogenetic tree, with no visible differences on the branches where egg or larval attendance evolved, namely *H. hampei* (median: 0.03), *P. cylindrus* (median: 0.04) and the ancestor of Scolytinae (median: 0.04), compared to all other weevil branches (medians: 0.00-0.04; Fig. S2). This indicates that there is no pervasive relaxation of selection across all tested orthologues on the branches where egg attendance and larval attendance evolved.

#### Relaxation and intensification of selection

To test if there are specific genes under relaxed or intensified selection on these three branches of interest compared to all other branches, we used RELAX (Wertheim et al. 2015) to analyse whether a change in strength of selection occurred with the emergence of parental care. Overall, there were more genes under relaxed (436) than under intensified (38) selection on the branches where egg and larval attendance evolved. The functional annotation of the top 10 genes under relaxed selection on the three branches where egg and larval attendance evolved (Table S3), revealed two clusters of functions: (1) gene expression regulation, more precisely - translation elongation, regulation of transcription, RNA processing (HELICc2) and pre-rRNA processing (rRNA adenine N(6)-methyltransferase family) and (2) neural plasticity, namely dystrobrevin binding protein 1 and KIF-1 binding protein. Two other proteins could be annotated with functions related to protein binding and signal transducer activity, while two more proteins have unknown functions (C-terminal to LisH motif and unknown). A lower number of genes were under intensified selection and had more diverse functions (Table S3). Three of the top 10 genes under intensified selection are related to gene expression regulation - specifically pre-mRNA splicing (LSM4 homolog), regulation of transcription and regulation of mRNA processing. Another protein, with the broad function “Cytoskeleton/chromatin conformation” might also be involved in transcriptional regulation through the changes in chromatin conformation. Two proteins are involved in protein transport, namely ADP-ribosylation factor family and Peptidase S24-like. Another two proteins are involved in electron transfer: namely ATP-dependent (S)-NAD(P)H-hydrate dehydratase and the mitochondrial ubiquinol-cytochrome C chaperone. Two further proteins play a role in carbohydrate metabolism: chitin metabolism and N-acetylgalactosaminyltransferase.

To test if parental care correlates with intensification or relaxation of selection across the phylogenetic tree, we ran RELAX for each single copy orthogroup with each branch set as the foreground branch. The proportion of orthogroups under significant relaxed selection was much higher in *P. cylindrus* (18.73%) than on all other branches, while *H. hampei* had the fourth highest value (4.86%, Fig. S4). On the other hand, the proportion of genes under intensified selection was very low in *P. cylindrus* (0.44%) and *H. hampei* (0.97%) the second and fourth lowest value, respectively (Fig. S5). Genes under relaxed selection on these focal branches are functionally enriched for the following GO terms: Regulation of gene expression, Regulation of cellular biosynthetic process, Regulation of biological process, Protein modification process, Regulation of transcription by RNA pol II, Cellular process, DNA-templated transcription.

These results are reflected in the median relaxation-intensification coefficient, log2(k), (Fig. 5 a). A positive log2(k) indicates intensification of selection, while a negative value indicates relaxation of selection. Both terminal branches with larval attendance have negative log2(k) values (*H. hampei* : −0.68, *P. cylindrus*: −0.78), but there are also species without any parental care with a negative log2(k). Se formally tested if there is a correlation between parental care and strength of selection, using a generalised linear mixed model (GLMM) as well as a Phylogenetic Generalised Least Squares model (PGLS).

**Figure 5:**
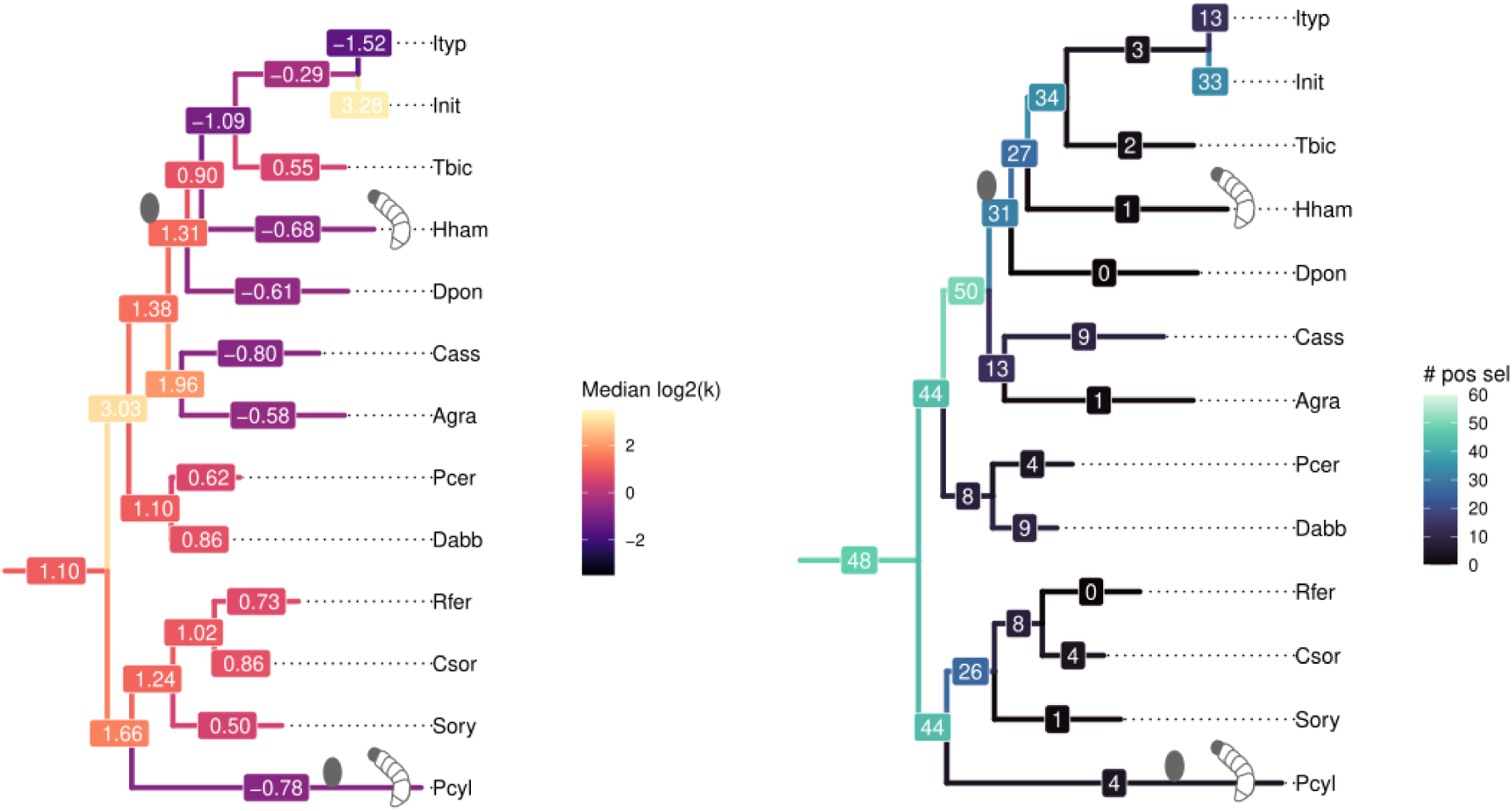
(a) Phylogenetic tree showing median log2(k) values for each branch. Eggs indicate where egg attendance evolved, while larvae indicate where larval attendance evolved. Positive values indicate intensification, negative values relaxation of selection. log2(k) was calculated for all single copy orthologues for each branch separately. For each branch, the median value of all significant (p.adjust) k values was then calculated. (b) Phylogenetic tree showing the number of genes under positive selection on each branch. For full species names please check Table S1.

Using all phenotypic characteristics shown in Fig. 1, namely pattern of care, parental care type, food type, genome size and ploidy as potential factors, we compared GLMMs of different size to each other. The intensification and relaxation parameter k was transformed using boxcox transformation to fit a normal distribution. The best model was *boxcox*(*k*) ∼ *parental care* + (1|*species*) + (1|*orthogroup*) with an AIC of 87880. The intensification and relaxation parameter (boxcox-transformed k) is lowest in larval attendance and highest in egg attendance. However, after multiple testing correction, the GLMM does not yield a significant difference in the relaxation parameter k between the different levels of parental care (no care, nest building, egg attendance, larval attendance). To take phylogenetic signals in the data into account, we performed a PGLS analysis for median k per branch (Fig. 5 a). We compared two models: *median k* ∼ *parental care, correlation* = *corBrownian*(1*, tree, form* =∼ *species*) and *median k* ∼ *parental care, correlation* = *corMartins*(1*, tree, form* =∼ *species*). The second model (corMartins) had a better fit according to logLik and AIC. According to this model, larval attendance has a significantly lower median intensification and relaxation parameter than nest building after multiple testing correction. This difference indicates that branches, on which larval attendance evolved, experienced relaxed selection compared to branches with nest building as the only parental care phenotype.

#### Positive selection

To test if positive selection plays a role in the evolution of egg or larval attendance, we used aBSREL (M. D. Smith et al. 2015) to calculate for each branch, how many genes showed signs of significant positive selection. On the branches where egg and larval attendance evolved, there were not more genes under positive selection than on other branches, namely 31 on the branch leading up to all Scolytinae, 4 on *P. cylindrus* and 1 on *H. hampei*, compared to 0 - 33 genes on all other weevil branches (0 - 50 genes on all branches, including ancestral ones, see Fig. 5 b). Furthermore, no genes were under convergent positive selection on more than one of the branches of interest. However, functional annotation (see Table S4) revealed genes related to transcriptional regulation to be under positive selection in *P. cylindrus* and the ancestor of Scolytinae, hinting at a possible role of changes in gene expression regulation in the evolution of egg attendance. Additionally, positive selection was observed in genes connected to microtubules and cytoskeletal structures in *H. hampei* and the Scolytinae ancestor. Interestingly, Moesin ezrin radixin and ATP-dependent microtubule severing protein, under positive selection in both *H. hampei* and the ancestor of Scolytinae, seems to be involved in axon targeting and axon regeneration. Together with the dystrobrevin binding protein 1 associated with neural plasticity under relaxed selection on all three branches of interest (Table S3) these findings indicate that neural development may be involved in the evolution of parental care. Genes under positive selection in the ancestor of all Scolytinae can be grouped into further functional categories, namely DNA and protein related processes, calcium homeostasis, microtubule related processes, respiratory electron chain, ion binding and others.

## Discussion

Subsociality, in which adults provide care for their offspring, is often regarded as a prerequisite for more socially complex phenotypes, such as eusociality, to evolve (Rehan & Toth 2015). The evolution of subsociality in insects is expected to coincide with changes in transcriptional regulation that influence caring behaviour, while adaptive evolutionary process are hypothesised to play a greater role in the evolution of more complex sociality (Rehan & Toth 2015; Pyenson & Rehan 2024). Furthermore, caring of dependent larvae may lead to a relaxation of selective forces on functions that are more important for survival in free-living, independent larvae (Mashoodh et al. 2023). In the weevil family, Curculionidae, various subsocial traits have evolved multiple times, while eusociality exists in a single species, making this an ideal clade for studying the molecular evolution of simple social phenotypes. We analysed genomes of 13 weevil species that span multiple combinations of care behaviours, including two independent origins of egg and larval attendance, to investigate the genomic mechanisms involved in the evolution of subsociality in weevils. We performed analyses of selection and gene family size evolution to test two main hypotheses: (1) the sheltering effects of parental care lead to relaxed selection on genes critical for larval survival; (2) the emergence of caring behaviour in weevils is associated with the adaptive evolution of transcriptional regulation.

### Sheltering of larvae leads to relaxed selection and gene losses in species with larval attendance

We expected parental care to act as a selective buffer in *P. cylindrus* and *H. hampei*, reducing the need for offspring to evolve traits that are necessary for independent larvae to survive in unprotected conditions. In support, we find over 400 genes to have experienced a significant relaxation of selection strength on the branches where egg and larval attendance evolved compared to only 38 genes with intensified selection. When comparing across the phylogeny, selection was significantly more relaxed in the two species with larval attendance, compared to species with just nest building, emphasising the sheltering effect of parental care of larvae. Mainly genes involved in transcriptional regulation and neural plasticity were under relaxed selection. The former may be associated with an overall reduction in protein synthesis linked to a reduced spectrum of behaviours in protected larvae. The latter may be related to changes in early brain development of dependent larvae. The inclusion of further genomes and transcriptomes representing varying combinations of social traits will help further test these hypotheses.

In further support of a link between relaxed selection and larval attendance, we found more convergently contracted gene families on branches where parental care evolved than on other branches, indicating some common gene losses among the species with parental care. Most of the functions of these genes can be divided into two groups, namely gene expression regulation and enzymatic activity, which may again be related to an overall reduction in the functional diversity of protected larvae.

Overall, across all single copy orthologues, we found genome-wide uniform strong purifying selection across the phylogenetic tree, including the species with parental care. This indicates that relaxed selection in weevils with larval attendance only affects specific functions, possibly those linked to larval survival, rather than being pervasive across the genome as a consequence of reduced effective population size, as has previously been suggested for eusocial species (Weyna & Romiguier 2021; C. Roux et al. 2024; Ewart et al. 2024).

### Adaptive changes in transcriptional regulation along with the emergence of egg and larval attendance

Adaptive evolution is hypothesised to play a greater role in the emergence of more advanced levels of insect sociality, than in the transitions from solitary to subsocial lifestyles, where mainly changes in transcriptional regulation are expected (Rehan & Toth 2015).

Accordingly, we find evidence of positive selection only in few genes (36) on the three branches where parental care of larvae or eggs evolved in our data set. However, consistent with expectations for the evolution of subsociality, many of these genes under positive selection are involved in transcriptional regulation. Specifically, in *P. cylindrus*, where egg and larval attendance evolved, we found that three out of four genes under positive selection had functions related to transcriptional regulation, namely orthogroups with sequence-specific DNA binding function, zinc finger protein 512B and a gene containing an HMG (high mobility group) box, all of which are involved in DNA binding during transcription.

In the ancestor of all Scolytinae species in this study, in which egg attendance evolved, we found 31 genes under positive selection, of which six have functions related to gene expression regulation and another eight genes have functions related to DNA and protein related processes. However, in *H. hampei*, where only larval attendance evolved, we found only one gene, Moesin ezrin radixin homolog, to be under positive selection. This gene plays a role in different processes, including mRNA export, axon targeting and connections of cytoskeletal structures and the plasma membrane (McCartney & Fehon 1996; Ruan et al. 2013; Kristó et al. 2017).

Importantly, genes with gene expression regulation function are also under intensified selection on the branches where parental care evolved. These genes were LSM4 homolog, likely involved in pre-mRNA splicing and a gene containing a domain found in Pit-Oct-Unc transcription factors likely involved in regulation of transcription, as well as a further gene involved in the regulation of mRNA processing. Additionally, a gene belonging to the actin family might play a role in transcriptional regulation through chromatin conformation changes. This suggests that transcriptional regulation plays an important role in the evolution of parental care.

## Conclusion

Our findings support the hypotheses that parental care contributes to sheltering, manifested as relaxed selection in a subset of genes, and that changes in parental care behaviours drive adaptive evolution in transcriptional regulation. Specifically, we observed gene loss or reduced functionality in genes associated with transcriptional regulation, suggesting relaxed selection in larvae whose behavioral repertoires may be reduced. Conversely, we detected signals of adaptive evolution or intensification in other regulatory genes, which may underpin behavioural innovations linked to parental care. These findings partially overlap with those linked to advanced eusocial phenotypes (Mikhailova et al. 2024), underscoring the need for finer representations of social phenotypes in genomic analyses to better capture the molecular processes at each stage of social evolution.

We propose that these dual trajectories, relaxed selection and adaptive evolution, reflect a fundamental tension in the evolution of subsociality: simplification of dependent behaviours and elaboration of caregiving traits. Future comparative genomic and transcriptomic studies, especially across diverse lineages exhibiting parental care, will be crucial for disentangling these dynamics and separating general evolutionary trends from species-specific phenomena.

## Materials and Methods

In this study, we included published genomes of 13 weevil species covering different parental care phenotypes, as well as two outgroups (the green lacewing *Chrysoperla carnea* Crowley et al. 2021 and the leaf-rolling weevil *Apoderus coryli* Crowley 2021). The two species with the most elaborate parental care phenotype are *Platypus cylindrus* Barclay, Vassiliades et al. 2024 (Platypodinae) and *Hypothenemus hampei* Navarro-Escalante et al. 2021 (Scolytinae) which independently evolved egg and larval attendance (Fig. 1). Four additional Scolytinae species with egg attendance, namely *Ips nitidus* Z. Wang et al. 2023, *Ips typographus* Powell et al. 2021, *Taphrorychus bicolor* Telfer & Badham 2024 and *Dendroctonus ponderosae* Keeling et al. 2022 suggest that egg attendance evolved in the ancestor of the five Scolytinae species in this study. Three other weevil species from two weevil tribes perform nest building, namely *Diaprepes abbreviatus* Sylvester et al. 2024, *Rhynchophorus ferrugineus* Dias et al. 2021 and *Sitophilus oryzae* Parisot et al. 2021. All other species deposit their eggs without providing any further care: *Anthonomus grandis grandis* A. K. Childers et al. 2021, *Ceutorhynchus assimilis* [Pest Genomics Initiative between Rothamsted Research, Bayer, and Syngenta], *Cosmopolites sordidus* Rodriguez Ruiz & Van Dam 2023, and *Polydrusus cervinus* Barclay, Natural History Museum Genome Acquisition Lab et al. 2023.

### Getting data

Genomes of 25 species (see Table S1) were downloaded from NCBI database in January 2024 and assembly quality was checked using BUSCO v 5.6.1 in genome mode with the insecta core set insecta_odb10 Manni et al. 2021. Of these species, 23 belong to the true weevils, two are leaf-rolling weevils, namely *Apoderus coryli* and *Cylas formicarius* and one species is the outgroup, namely *Chrysoperla carnea*. Genomes with BUSCO completeness scores higher than 97% were re-annotated.

### Re-annotation

For the comparable annotation of all genomes, we followed the FastTE Bell et al. 2022 pipeline for the repeat annotation and masking. Briefly, we used EDTA v2.1.3 Ou et al. 2019 and DeepTE Yan et al. 2020 for *de novo* TE annotation and classification, followed by repeat masking using RepeatMasker Smit et al. 2015. The repeat-masked genomes were then trimmed using Trimmomatic Bolger et al. 2014 to avoid adapter contamination. BRAKER 3.0.2 Brůna, Hoff et al. 2021; Hoff, Lomsadze et al. 2019; Hoff, Lange et al. 2016; Stanke, Schöffmann et al. 2006; Stanke, Diekhans et al. 2008; Brůna, Lomsadze et al. 2020; Lomsadze et al. 2005; Buchfink, Xie et al. 2015; Gotoh 2008; Iwata & Gotoh 2012 was used for genome annotation using the OrthoDB v11 eukaryota protein set. The quality of the genome annotation was assessed using BUSCO Manni et al. 2021 in protein mode and DOGMA Dohmen et al. 2016. Using a BUSCO completeness score of 95% as a cut-off, 15 species remained in this study (see bold names in Table S1).

Coding sequences (cds) were extracted using gffread from cufflinks 2.2.1 Trapnell et al. 2010 and protein domains were annotated using PfamScan Finn et al. 2014.

### Orthology, alignment and phylogenetic tree construction

SequenceFairies seqCheck was used to remove stop codons from the re-annotated proteome file and isoformCleaner to remove all but the longest isoform Kemena 2024. Orthogroups were inferred using OrthoFinder v.2.5.4 Emms & Kelly 2019 with default settings. Single copy orthologues were used for selection analyses, while multicopy orthologues were used for gene family size analyses. Proteome sequences were aligned with PRANK v.170427 Löytynoja 2014. Subsequently, a phylogenetic tree was constructed using iqtree Minh et al. 2020 and rooted with iTol v5 Letunic & Bork 2021.

### Gene family size changes

A preliminary analysis showed that many of the expanded and contracted gene families had annotations that contained transposable elements (TEs). To ensure that these elements with large copy number variations do not interfere with the analysis of gene family size changes, we performed an additional filtering step on all proteomes. Specifically, we removed all genes containing the following Pfam domains: PF00075, PF00078, PF00665, PF02925, PF02992, PF03184, PF03221, PF03732, PF04687, PF05699, PF05840, PF05970, PF07727, PF08283, PF08284, PF10551, PF13358, PF13359, PF13456, PF13837, PF13976, PF14214, PF14223, PF14529, taken from Min & Choi 2019. We also ran transposonPSI (Brian J. Haas, TransposonPSI, 2007-2011 <http://transposonpsi.sourceforge.net>) and combined both lists of genes. We removed these genes from the proteomes, and subsequently reran OrthoFinder as described above. Gene familiy size changes across the phylogenetic tree were then analysed using CAFE v5.0. Mendes et al. 2021. The rooted phylogenetic tree was transformed to an ultrametric tree using the make_ultrametric.py script that comes with OrthoFinder. The clade_and_size_filter.py script was used to subset small and large gene families, subsequent analyses were performed on the small gene families. We ran CAFE v5.0. multiple times with different gamma values to see which gamma value has the highest final likelihood. In this case, gamma = 1 had the highest likelihood score. The testing of different lambda values didn’t make sense biologically, so we chose lambda = 1. The values obtained were then used for analysis of the large gene families, but the analysis failed. Results were plotted using CafePlotter (https://github.com/moshi4/CafePlotter). A custom R script was used to analyse convergent contractions and expansions of gene families on 3 branches of interest. Functional annotation of convergently expanded and contracted gene families was performed using eggnog-mapper version 2.1.12 Cantalapiedra et al. 2021; Huerta-Cepas et al. 2019 with eggNOG DB version: 5.0.2, diamond version 2.1.10 Buchfink, Reuter et al. 2021.

### Selection analyses

For the selection analyses, the .codingseq files from the re-annotation with BRAKER3 was used. Coding sequences for each single copy orthogroup were extracted using Sequence-Fairies seqExtract Kemena 2024 and aligned based on the proteome files using pal2nal (v14) Suyama et al. 2006. The coding sequence alignment was trimmed using Gblocks (0.91) Castresana 2000 with the parameters -t=c -b5=h. A custom python script was then used to format Gblock output files as in fasta format.

#### Estimation of dN/dS values

The dN/dS values (ratio of the number of non-synonymous substitutions per non-synonymous site to number of synonymous substitutions per synonymous site), as an indicator for selective pressure, were calculated for each branch in the species tree and for each single copy orthologue. The calculation was performed using the free-ratios model implemented by CodeML in PAML suite (v. 4.9h) Yang 2007. dN/dS values were extracted using a custom python script. Analyses were performed in R R Core Team 2023 using RStudio Posit team 2023. A custom R script was used to plot median dN/dS per branch on the phylogenetic tree using the packages ggtree Yu, D. Smith et al. 2017; Yu, Lam et al. 2018; Yu 2020; Xu et al. 2022; Yu 2022, ggplot2 Wickham 2016, viridis Garnier et al. 2024, treeio L.-G. Wang et al. 2020.

#### Test of selection

To test if parental care is associated with a change in selection regime (relaxation or intensification of selection), RELAX Wertheim et al. 2015 from the HyPhy suite (v. 2.5.6) Pond et al. 2005 was used with default options. RELAX was run for each single copy orthogroup with the branches where parental care evolved, namely ancestor of Scolytinae, *H. hampei* and *P. cylindrus*, set as foreground branches. Subsequently p-values were corrected for multiple testing using the false discovery rate (FDR). To compare the functions of the top 10 genes under relaxed/intesified selection on the branches of interest, genes were functionally annotated using eggnog-mapper version 2.1.12 with eggNOG DB version: 5.0.2, diamond version 2.1.10. The distribution of k (intensification and relaxation parameter) was plotted using a custom R script with the packages ggplot2 and dplyr Wickham et al. 2023.

To generate data for linear modeling, we ran RELAX 27 times for each single copy orthogroup, each time with a different branch as foreground branch. A custom python script was used to extract genes under relaxed/intensified selection. The median-k-tree was plotted using a custom R script using the packages ggtree, ggplot2, viridis, dplyr and ape Paradis & Schliep 2019. To test if the observed differences in median k are statistically significant, we used a Phylogenetic Generalized Least Square (PGLS) model with Brownian Correlation Structure as well as Martins and Hansen’s (1997) covariance structure. The custom R script for these analyses made use of the libraries ape, nlme J. Pinheiro et al. 2025; J. C. Pinheiro & Bates 2000, stargazer Hlavac 2022, dplyr, emmeans Lenth 2025 and treeio.

To test for positive selection of genes in the phylogeny, aBSREL M. D. Smith et al. 2015 from the HyPhy suite (v.2.5.6) was used with default options. To test if the branches where parental care evolved are subject to episodic selection, we ran aBSREL with a priori selecting the foreground branches. Then, to perform an exploratory analysis, we ran aBSREL for each single copy orthogroup without foreground branches. A custom bash script was used to count the number of genes under positive selection per branch. Functional annotation was performed for the genes of interest as described above and a tree showing number of genes under positive selection was plotted using a custom R script using the same packages as for the median-k-tree.

## Data Availability

Genome assemblies were publicly available at NCBI (accession numbers see Table S1). Genome annotations, namely coding sequences (cds) and proteomes produced in this study are available at https://doi.org/10.5281/zenodo.16760856.

## Acknowledgements

SR is supported by the DFG Grant 503320462 to MH.

## Supplements

### Tables

**Table S1:**
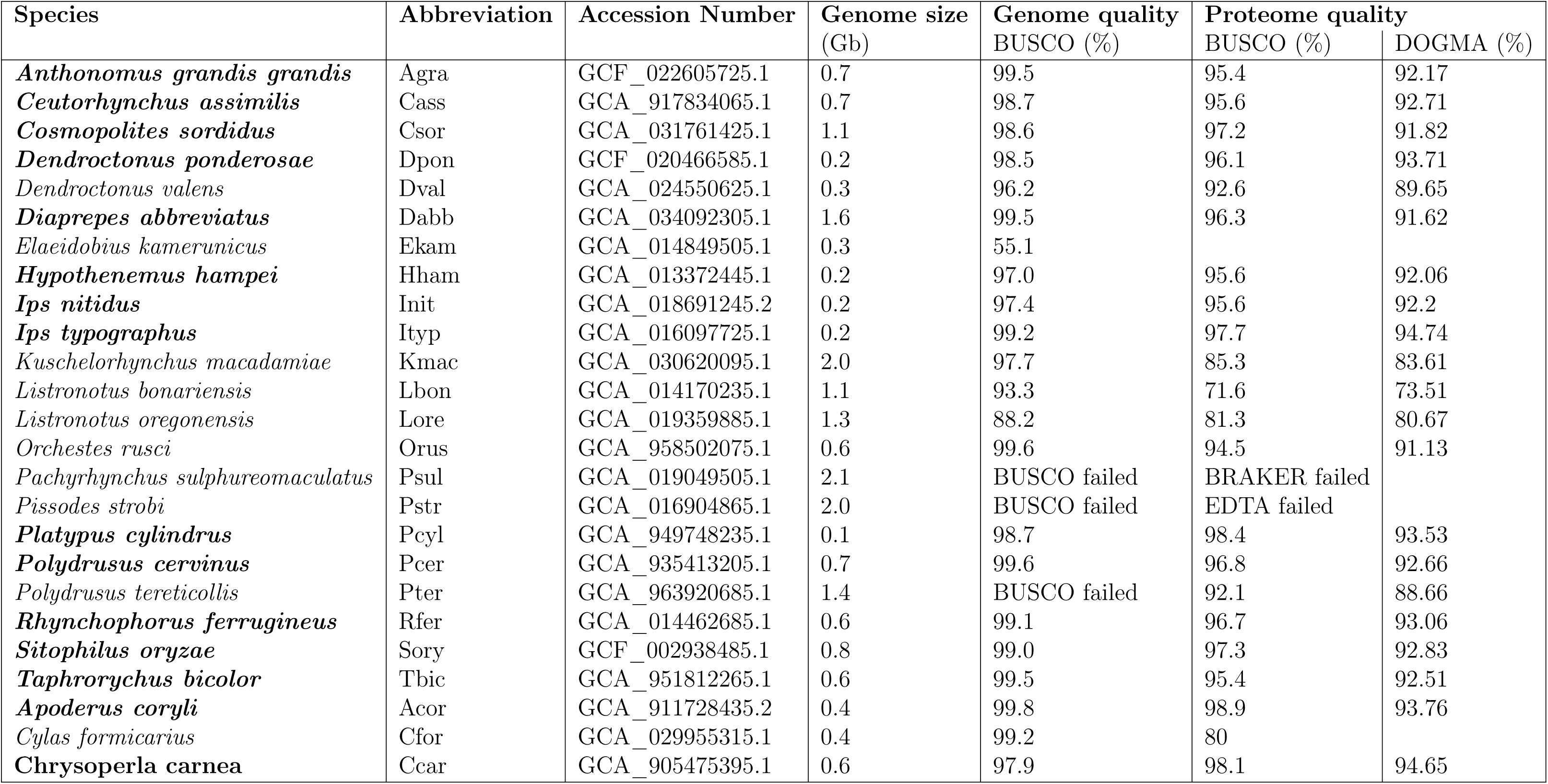
Quality assessment of genomes downloaded from NCBI and re-annotated proteomes using BUSCO and DOGMA. Species with BUSCO genome completeness score of >97% and BUSCO proteome completeness score of >95% were included in this study.

**Table S2:**
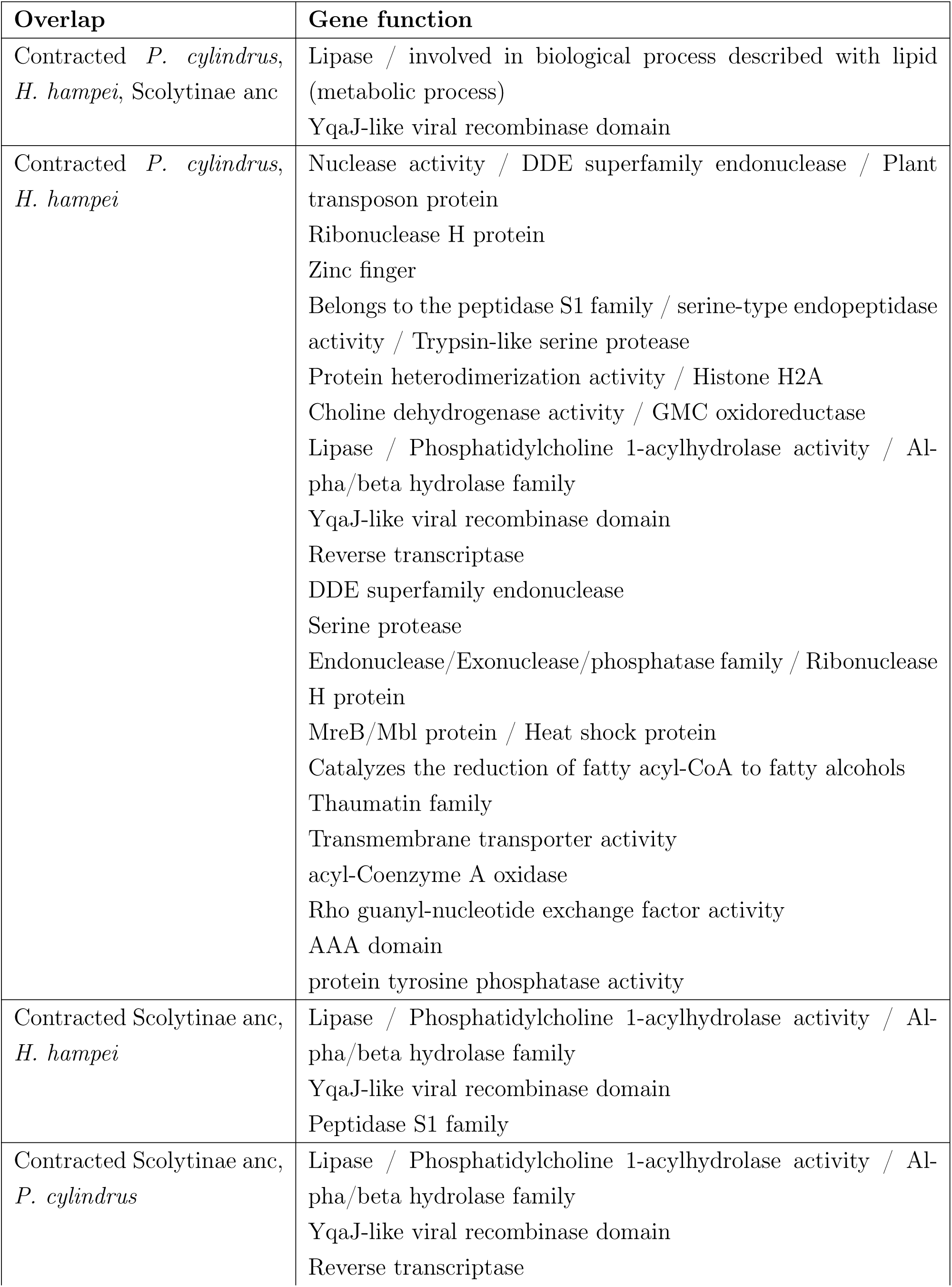
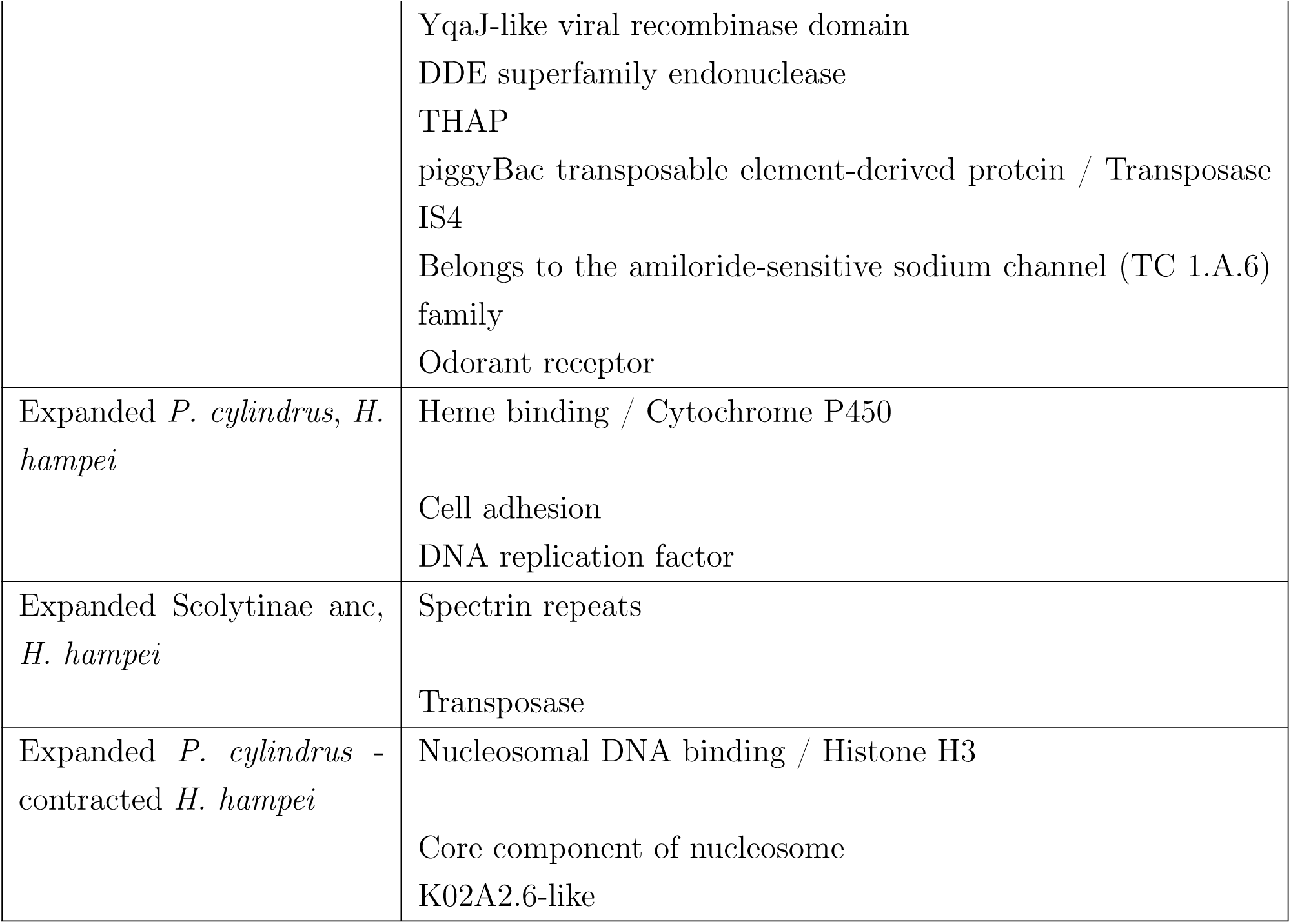
Functions of genes convergently expanded or contracted in different species (Overlap). Methods: CAFE5, cafeplotter, R, eggnog. Just showing the most common assigned function for each OG. Number of gene functions does not correspond to number of genes in overlaps, as not all genes were annotated by eggnog. OGs present in more than one overlap are shown in each overlap.

**Table S3:**
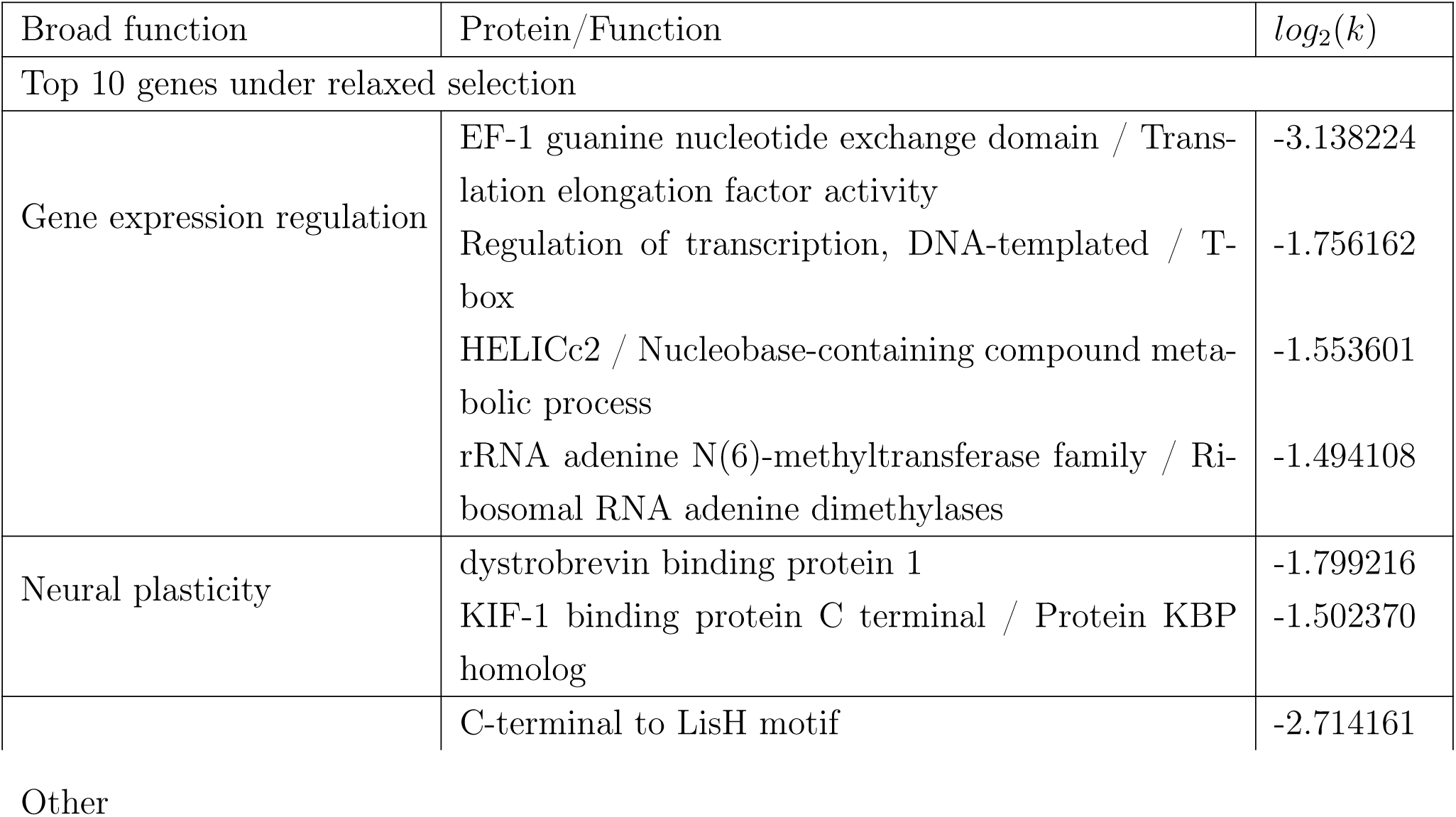
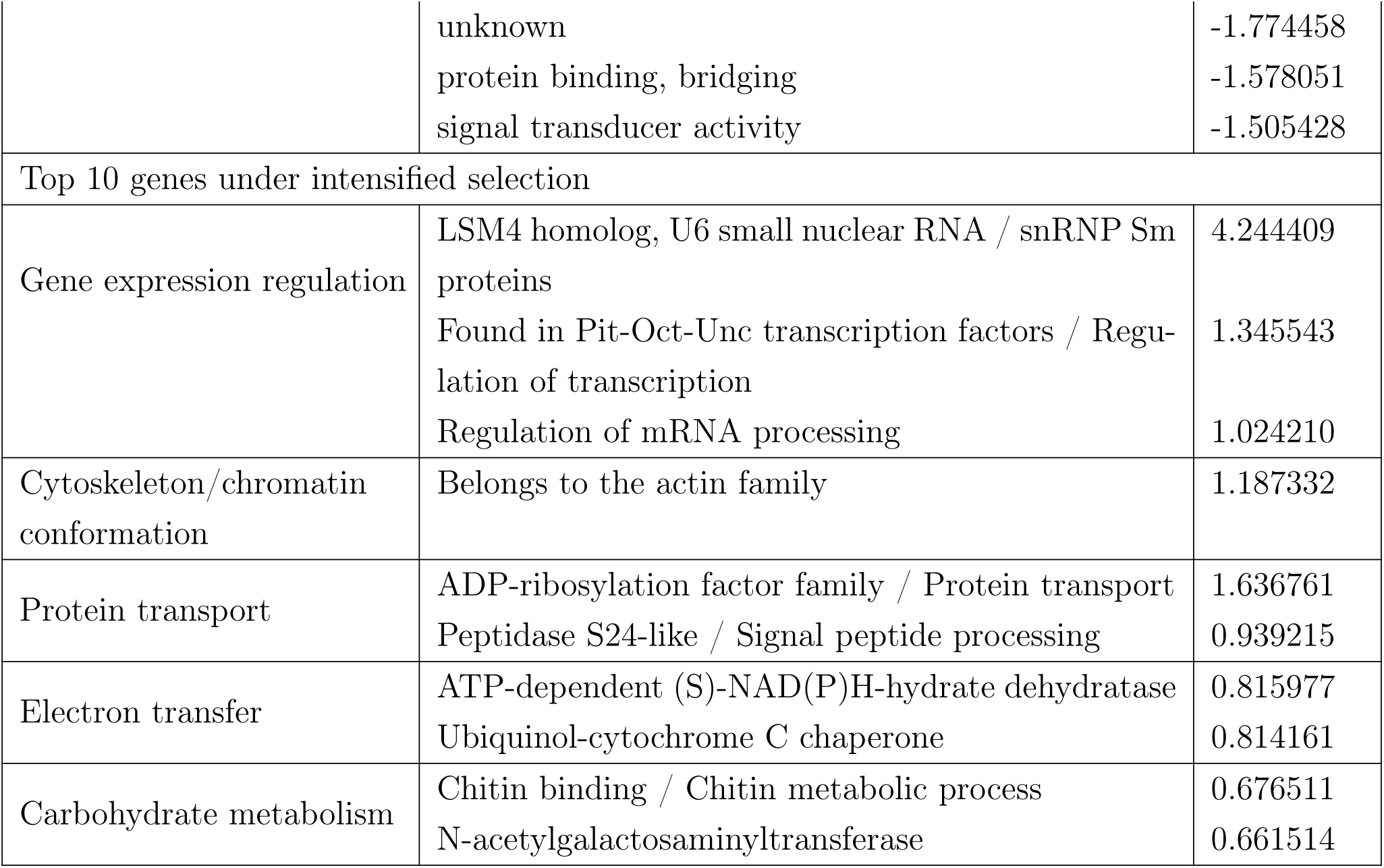
Functions of the top 10 genes under relaxed and under intensified selection in the ancestor of Scolytinae, *H. hampei* and *P. cylindrus*. Broad function contains a manual grouping of proteins with similar functions, Protein/Function contains a shortened output of eggnog functional annotation. *log*_2_(*k*) shows the intensification and relaxation parameter from the RELAX analysis, positive values indicate intensification of selection, negative values indicate relaxation of selection.

**Table S4:**
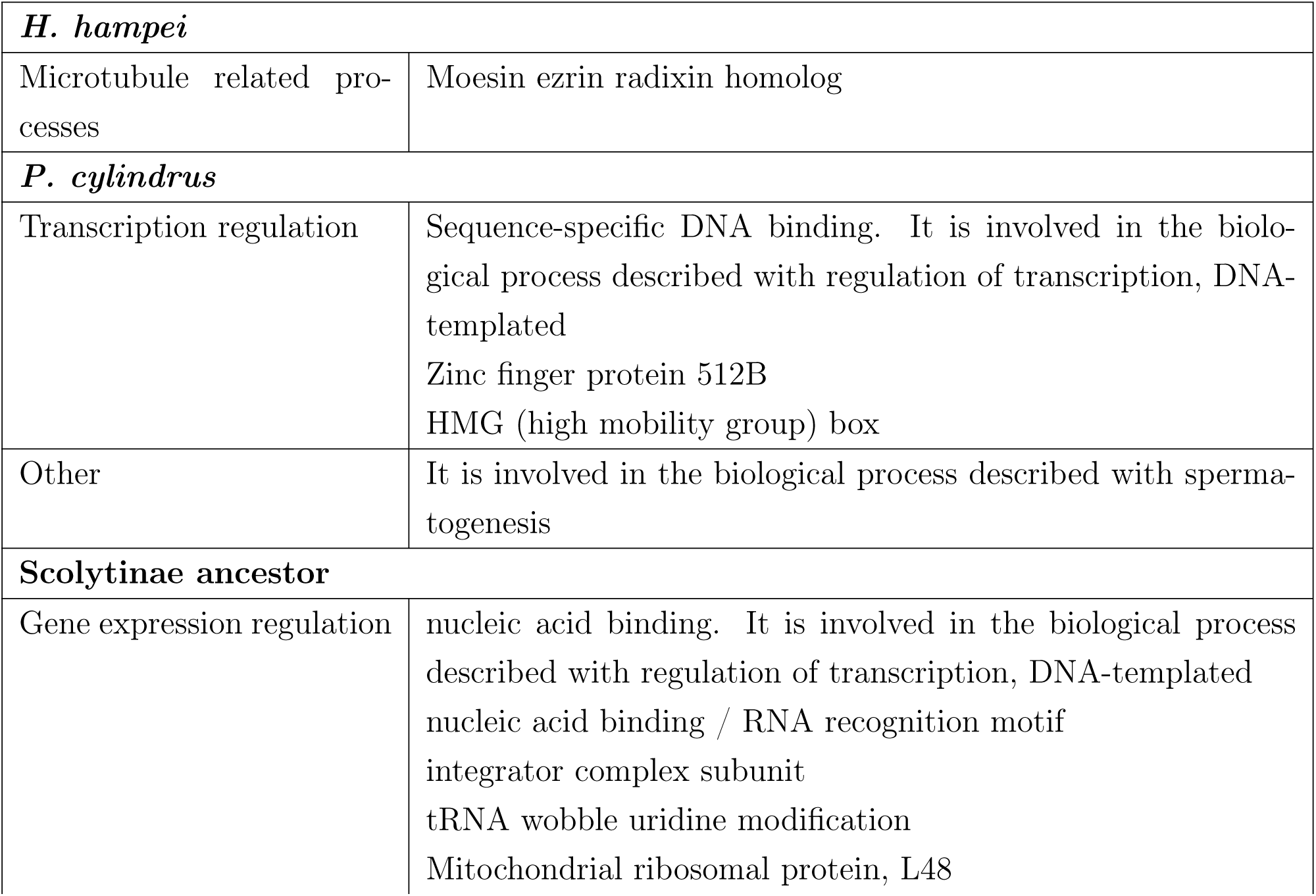
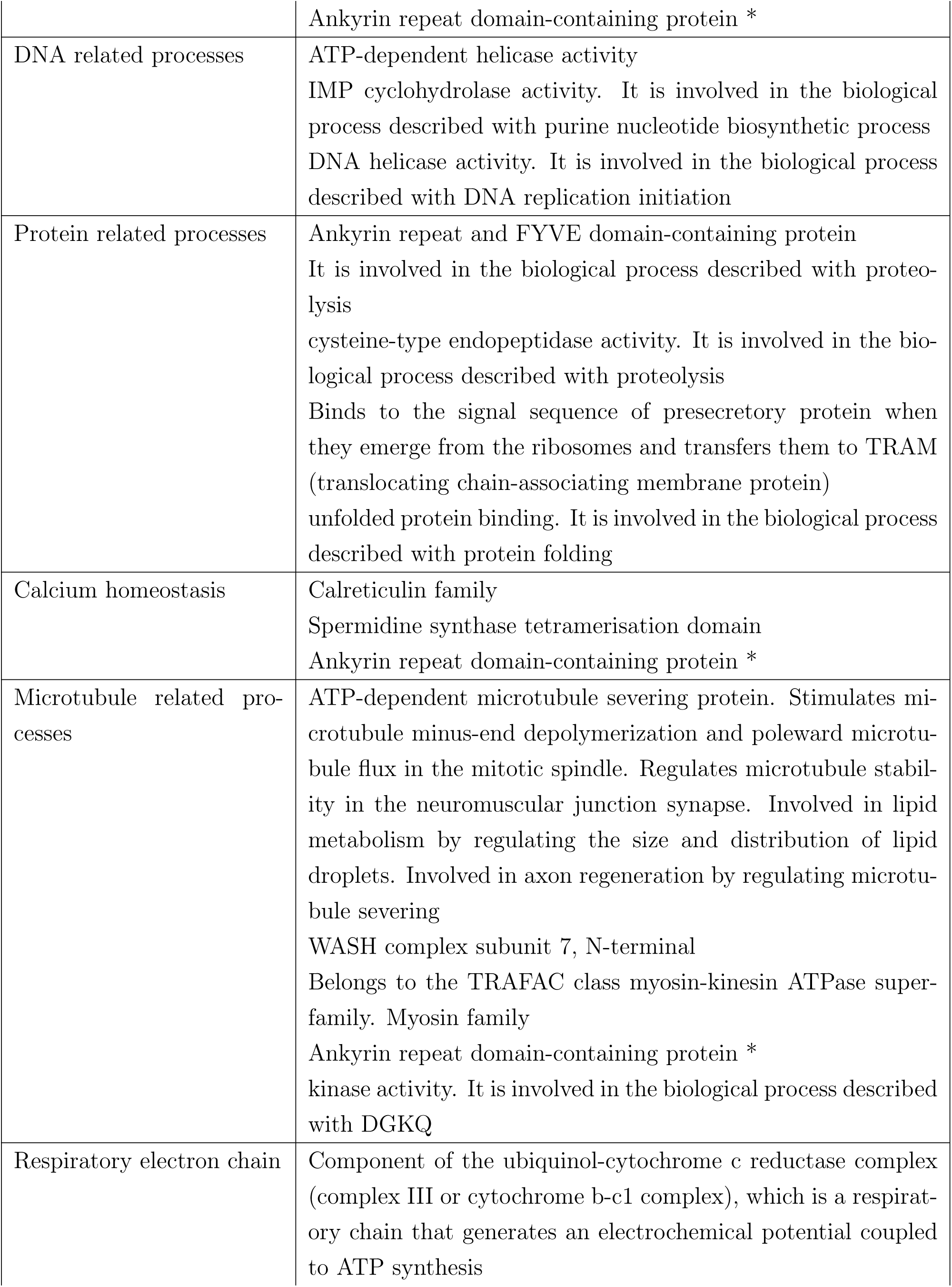
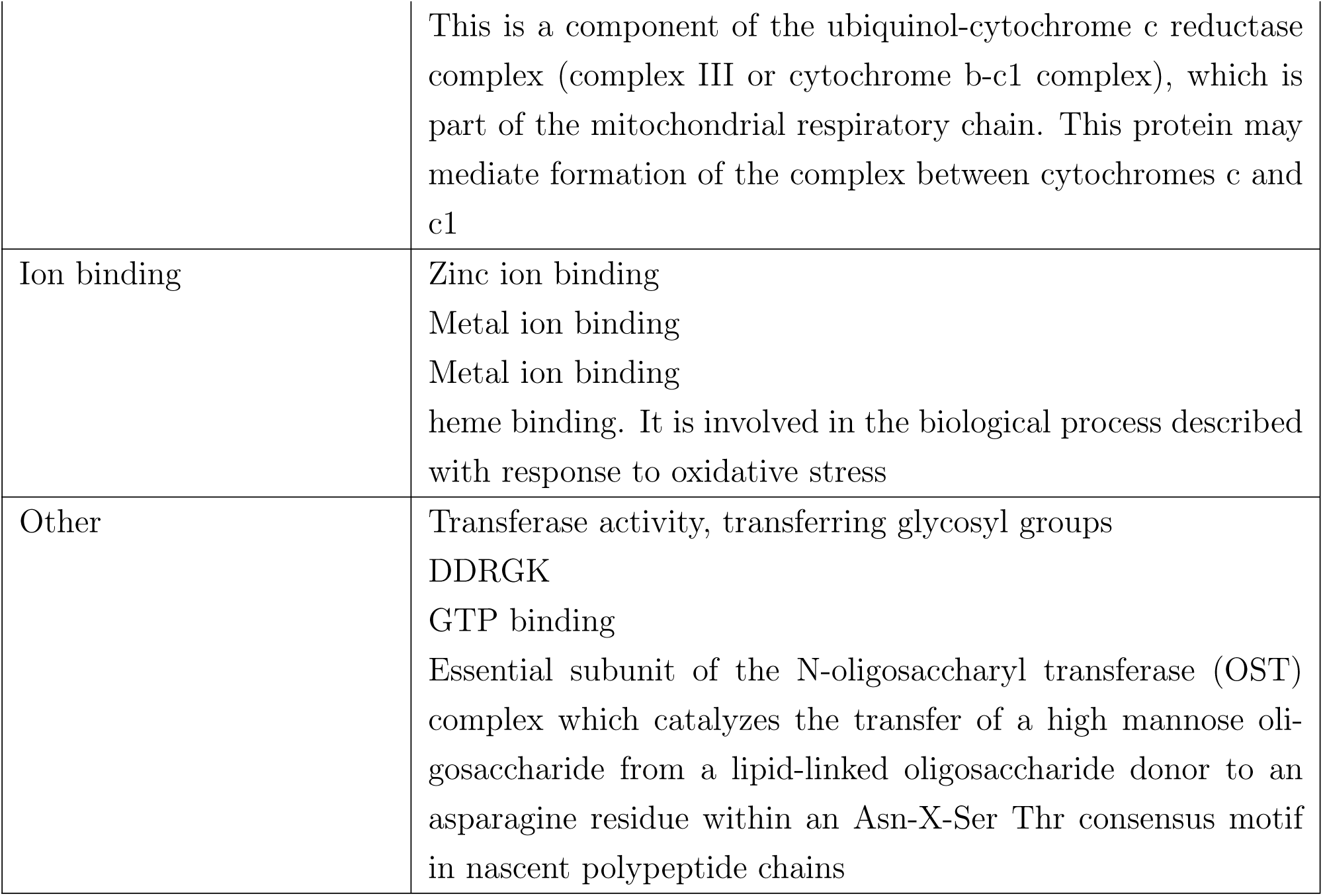
Functions of genes under positive selection in the ancestor of Scolytinae, *H. hampei* and *P. cylindrus*. *Ankyrin repeat domain-containing protein was only found to be under positive selection once, but could be grouped with different functional groups.

|  |  |
| --- | --- |
| ***H. hampei*** |  |
| Microtubule related processes | Moesin ezrin radixin homolog |
| ***P. cylindrus*** |  |
| Transcription regulation | Sequence-specific DNA binding. It is involved in the biological process described with regulation of transcription, DNA-templated<br>Zinc finger protein 512B<br>HMG (high mobility group) box |
| Other | It is involved in the biological process described with spermatogenesis |
| **Scolytinae ancestor** |  |
| Gene expression regulation | nucleic acid binding. It is involved in the biological process described with regulation of transcription, DNA-templated<br>nucleic acid binding / RNA recognition motif<br>integrator complex subunit<br>tRNA wobble uridine modification<br>Mitochondrial ribosomal protein, L48 |
|  | Ankyrin repeat domain-containing protein * |
| DNA related processes | ATP-dependent helicase activity<br>IMP cyclohydrolase activity. It is involved in the biological process described with purine nucleotide biosynthetic process<br>DNA helicase activity. It is involved in the biological process described with DNA replication initiation |
| Protein related processes | Ankyrin repeat and FYVE domain-containing protein<br>It is involved in the biological process described with proteolysis<br>cysteine-type endopeptidase activity. It is involved in the biological process described with proteolysis<br>Binds to the signal sequence of presecretory protein when they emerge from the ribosomes and transfers them to TRAM (translocating chain-associating membrane protein)<br>unfolded protein binding. It is involved in the biological process described with protein folding |
| Calcium homeostasis | Calreticulin family<br>Spermidine synthase tetramerisation domain<br>Ankyrin repeat domain-containing protein * |
| Microtubule related processes | ATP-dependent microtubule severing protein. Stimulates microtubule minus-end depolymerization and poleward microtubule flux in the mitotic spindle. Regulates microtubule stability in the neuromuscular junction synapse. Involved in lipid metabolism by regulating the size and distribution of lipid droplets. Involved in axon regeneration by regulating microtubule severing<br>WASH complex subunit 7, N-terminal<br>Belongs to the TRAFAC class myosin-kinesin ATPase superfamily. Myosin family<br>Ankyrin repeat domain-containing protein *<br>kinase activity. It is involved in the biological process described with DGKQ |
| Respiratory electron chain | Component of the ubiquinol-cytochrome c reductase complex (complex III or cytochrome b-c1 complex), which is a respiratory chain that generates an electrochemical potential coupled to ATP synthesis |
|  | This is a component of the ubiquinol-cytochrome c reductase complex (complex III or cytochrome b-c1 complex), which is part of the mitochondrial respiratory chain. This protein may mediate formation of the complex between cytochromes c and c1 |
| Ion binding | Zinc ion binding<br>Metal ion binding<br>Metal ion binding<br>heme binding. It is involved in the biological process described with response to oxidative stress |
| Other | Transferase activity, transferring glycosyl groups<br>DDRGK<br>GTP binding<br>Essential subunit of the N-oligosaccharyl transferase (OST) complex which catalyzes the transfer of a high mannose oligosaccharide from a lipid-linked oligosaccharide donor to an asparagine residue within an Asn-X-Ser Thr consensus motif in nascent polypeptide chains |

### Figures

**Figure S1:**
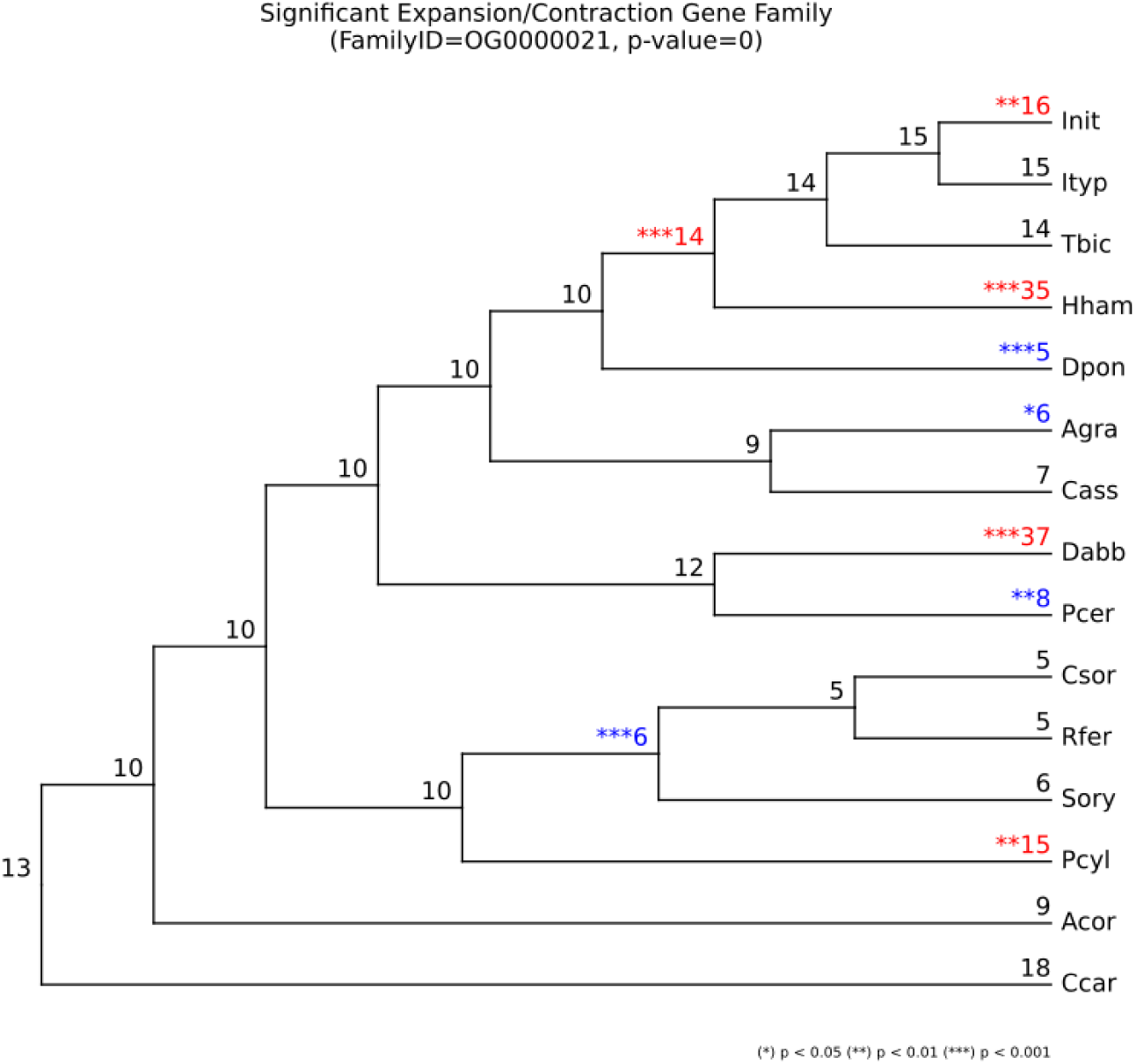
Gene family size evolution of the orthogroup annotated as Heme binding / Cytochrome P450. Red numbers indicate significant expansions, blue numbers indicate significant contractions. This gene family is significantly expanded in both, *H. hampei* (35 gene copies) and *P. cylindrus* (15 gene copies). However, it is also significantly expanded in two other species and one ancestral branch. For full species names please check Table S1.

**Figure S2:**
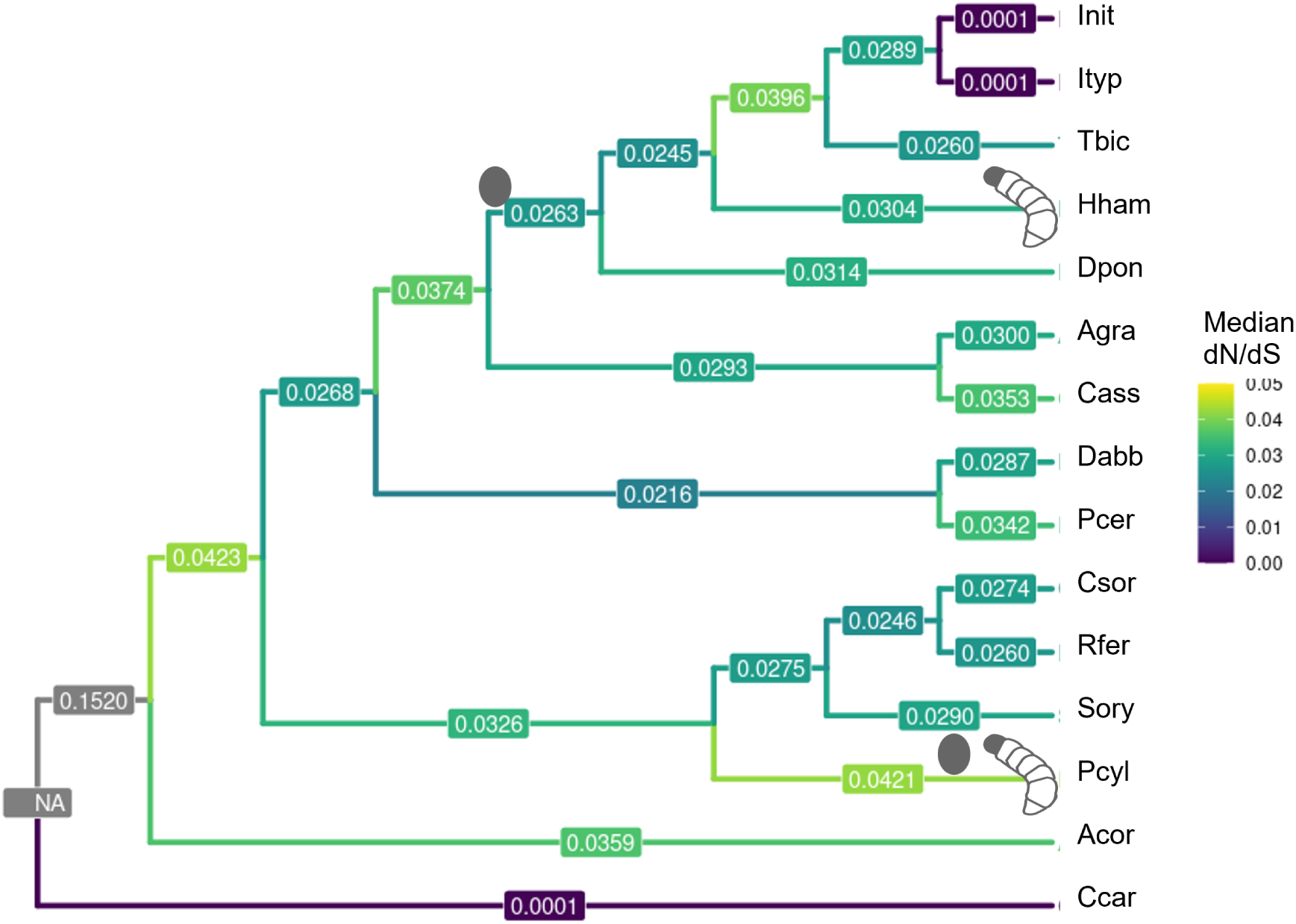
Median dN/dS tree. dN/dS was calculated for each single copy orthologue using CodeML, the median per branch was then calculated and plotted. A dN/dS value below 1 indicates purifying selection, above 1 positive selection. Altogether, selection is strongly purifying across the tree without large differences between the branches.

**Figure S3:**
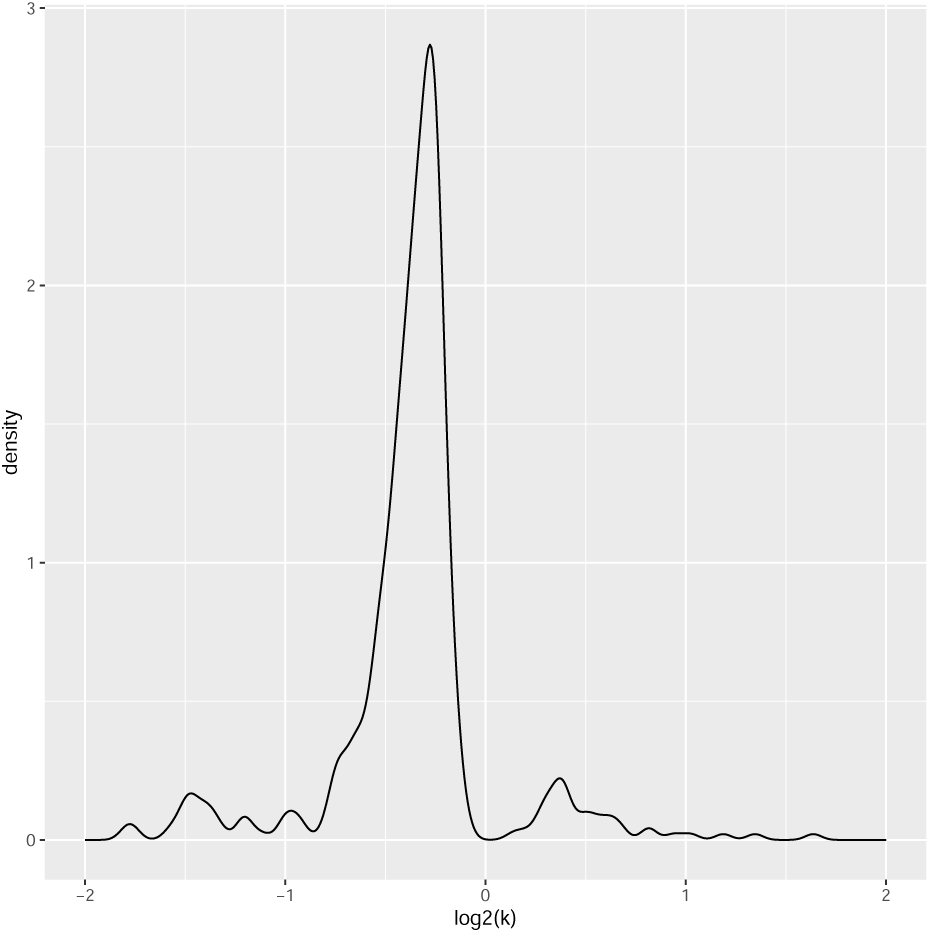
Relax: distribution of log2(k): Below zero - relaxation, above zero - intensification. Here, the change of intensity of selection in the three focal branches (*H. hampei*, *P. cylindrus*, Scolytinae anc) compared to all background branches is analysed

**Figure S4:**
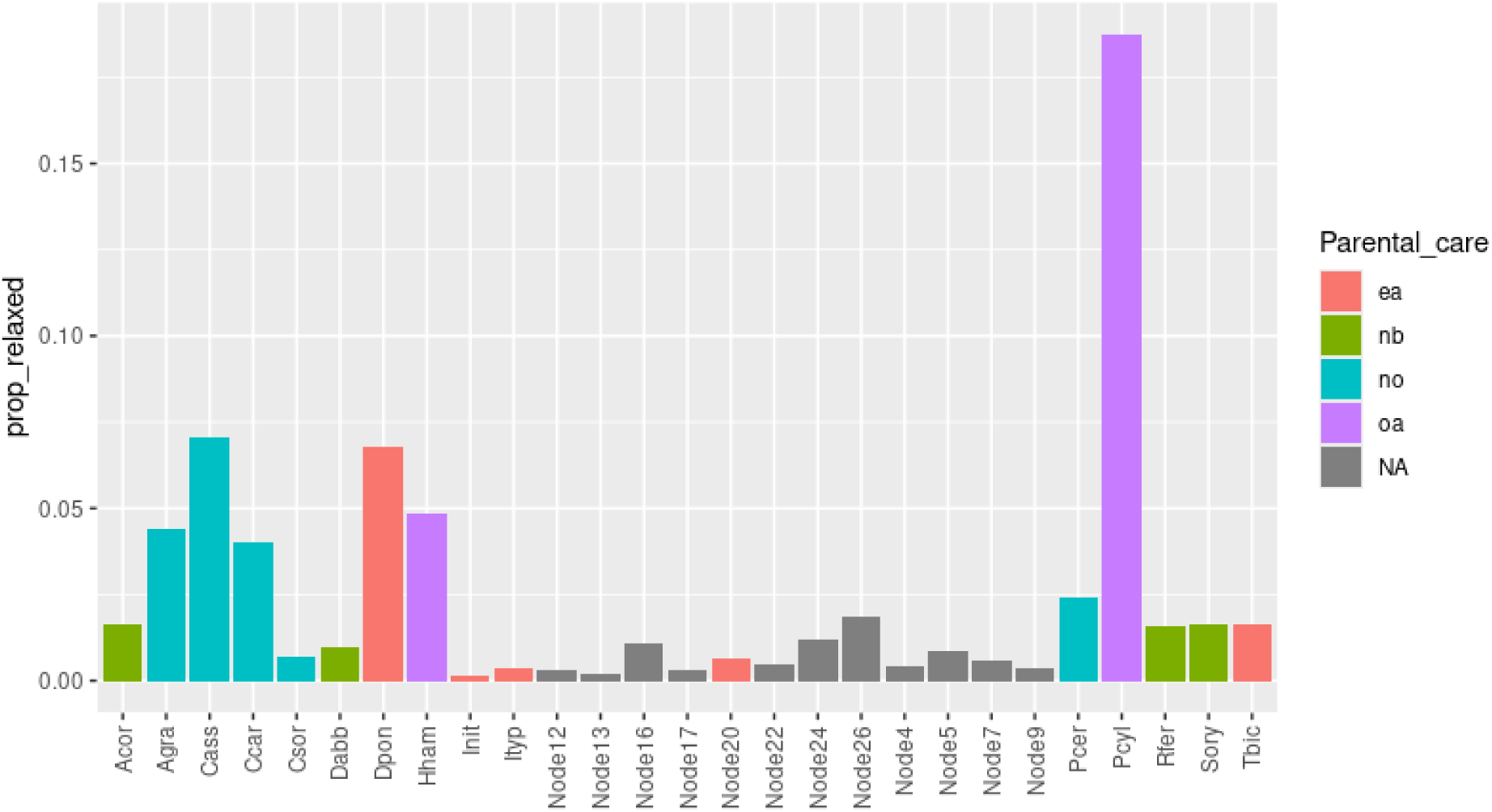
Proportion of orthogroups under relaxed selection on each branch. log2(k) was calculated for all single copy orthologues for each branch separately. For each branch, the median value of all significant (p.adjust) k values was then calculated.

**Figure S5:**
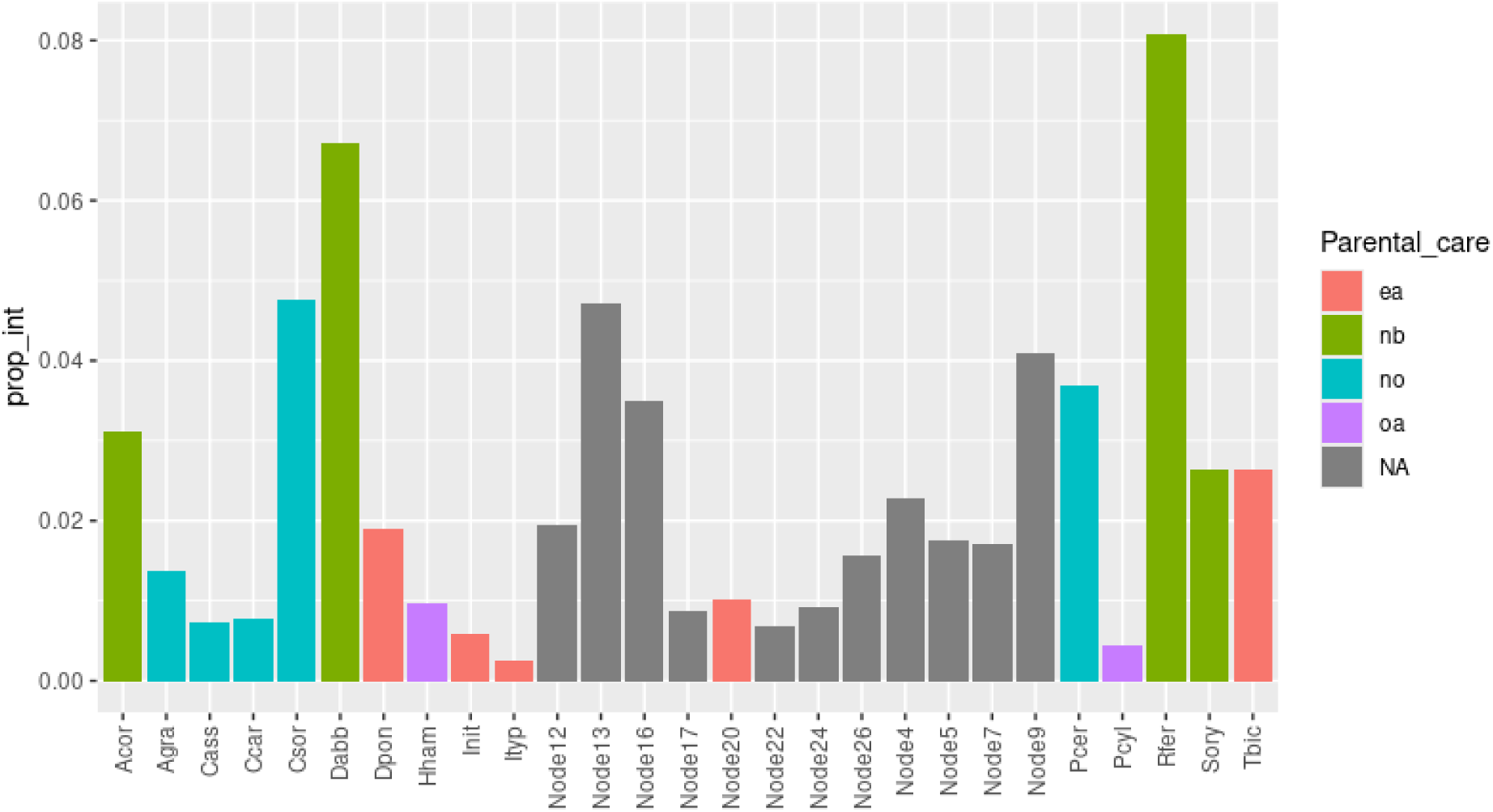
Proportion of orthogroups under intensified selection on each branch. log2(k) was calculated for all single copy orthologues for each branch separately. For each branch, the median value of all significant (p.adjust) k values was then calculated.

